# C1q from C1q^+^ tumor-associated myeloid cells promotes resistance to T-cell engagers and CAR T-cells and is induced by LIF and glucocorticoids

**DOI:** 10.64898/2026.09.15.751784

**Authors:** María López-Vázquez, Sergio Espinosa-Gil, Raffaella Iurlaro, Laura Carrillo-Bosch, Ariadna Grinyó-Escuer, Darío Solís-Sayago, Alexandra Arias, Isabel Cuartas, Almudena Neva-Alejo, Laia Cuesta-Casanovas, Ester Bonfill-Teixidor, Cayetano Galera-Martínez, César Merino-Allona, María Duarte-León, Irepan Salvador, Pablo Clavero, Marta Giménez-Alejandre, Holger Heyn, Josep González, Estela Pineda, Sonia Guedan, Francisco Martínez-Ricarte, Joan Seoane

## Abstract

Immunotherapies, particularly T-cell engagers (TCEs) and CAR T-cells, have shown limited efficacy in solid tumors, partly due to an immunosuppressive tumor microenvironment (TME). However, the molecular mechanisms by which the TME impairs immunotherapy remain poorly understood. Here, we found that C1q generated by C1q⁺ tumor-associated myeloid cells (TAMs) plays a fundamental role in shaping the immunosuppressive TME in glioblastoma (GBM), one of the most aggressive tumors. C1q suppressed T-cell activation and impaired the activity of T-cell engagers (TCEs) and CAR T-cells. Genetic ablation of *C1qa* improved anti-tumor responses to TCEs and CAR T-cells. We used innovative patient-derived tumor tissue cultures (PDTTCs), which preserve an intact TME, from 19 GBM patients and identified the LIF cytokine as the main inducer of C1q. Moreover, we discovered that glucocorticoids cooperate with LIF to induce C1q. The identified C1q⁺ TAM signature overlapped with an anti-LIF gene signature, was associated with poor prognosis, and was enriched in mesenchymal GBMs with *NF1* mutations. The blockade of LIF using an anti-LIF neutralizing antibody decreased the presence of C1q^+^ TAMs, and we found that the regulation of C1q by anti-LIF is conserved between human and mouse. Using the *C1qa*^⁻/⁻^ GBM mouse model, we showed that C1q mediates the anti-tumor immune response induced by LIF blockade. Our findings identify C1q⁺ TAMs as key immunosuppressive players in GBM, impairing CAR T-cells and TCE activity, and position them as therapeutic targets to improve immunotherapy responses in this devastating disease.

---

Immunotherapies, particularly T-cell based therapies (TCTs, i.e., T-cell engagers (TCEs) and CAR T-cells), have failed to provide benefit for patients with solid tumors including glioblastoma (GBM)^1^. GBM is the most common aggressive primary brain tumor in adults, a dismal disease with a clear medical unmet need^2,3^. TCTs have shown remarkable results in hematological malignancies^4–8^. However, their development in solid tumors has been hampered by the negative impact of the tumor microenvironment (TME)^9,10^. Still, the molecular players hindering the effectiveness of TCTs in solid tumors have not yet been fully elucidated.

Tumor-associated myeloid cells (TAMs) are some of the most prevalent non-malignant cell type in GBM, reaching up to 30% of all cells in a tumor^11^. Recent studies have begun to uncover the diversity and heterogeneity of TAM cell states^12,13^. However, the molecular determinants of specific TAM phenotypes and their immunosuppressive function are still unknown.

One of the impediments to study the TME and discern TAM function in human cancer relates to challenges with experimental modeling of the human TME. Tumor organoids do not contain immune cells, such as TAMs, and patient-derived xenograft (PDX) models do not recapitulate the patient’s myeloid compartment^14^. In order to circumvent these issues, we developed *ex vivo* tumor tissue cultures, referred to as patient-derived tumor tissue cultures (PDTTCs), which provide a fully human platform that accurately recapitulates the patient’s tumor including the TME and its immune cells. This model is described in (Iurlaro et al. co-submitted).

The C1q protein complex encoded by three genes, *C1QA*, *C1QB*, and *C1QC*, is a member of the complement system primarily expressed extra-hepatically in myeloid cells that can exert a tolerogenic function by inhibiting T-cells^15,16^. Several reports have shown that TAMs expressing C1q (C1q^+^ TAMs) are mediators of immunosuppression in cancer^17–19^. However, the molecular mechanisms that control their presence in tumors and their impact on TCEs and CAR T-cells remain unknown.

We and others have identified the LIF cytokine as a therapeutic target in cancer that can signal through TAMs^20–24^. On the other hand, glucocorticoids (i.e., dexamethasone) have long been employed as a first-line therapeutical strategy in the management of GBM induced and surgery associated edema^25,26^. However, the molecular impact of glucocorticoids on immunotherapies is still under study.

Using patient samples, patient-derived models, genetically modified and syngeneic mouse models, we found that C1q secreted by TAMs inhibits T-cell activity and impairs the response to TCEs and CAR T-cells. Moreover, we discovered that C1q is induced by the cooperation of LIF and glucocorticoids, promoting an immunosuppressive TME.

Thus, we identified C1q^+^ TAMs, regulated by both LIF and glucocorticoids, as therapeutic targets to overcome the immunosuppressive TME and enable responses to TCEs and CAR T-cells.

## Results

### C1q inhibits the response to TCEs and CAR T-cells

We recently identified C1q^+^ expressing TAMs in GBM (Iurlaro et al. co-submitted). We analyzed the single-cell RNA sequencing data (scRNAseq) and identified a C1q^+^ cell cluster exclusively present in the myeloid compartment defined by a C1q^+^ TAM gene signature (Fig. 1a, 1b and S1).

**Fig. 1.**
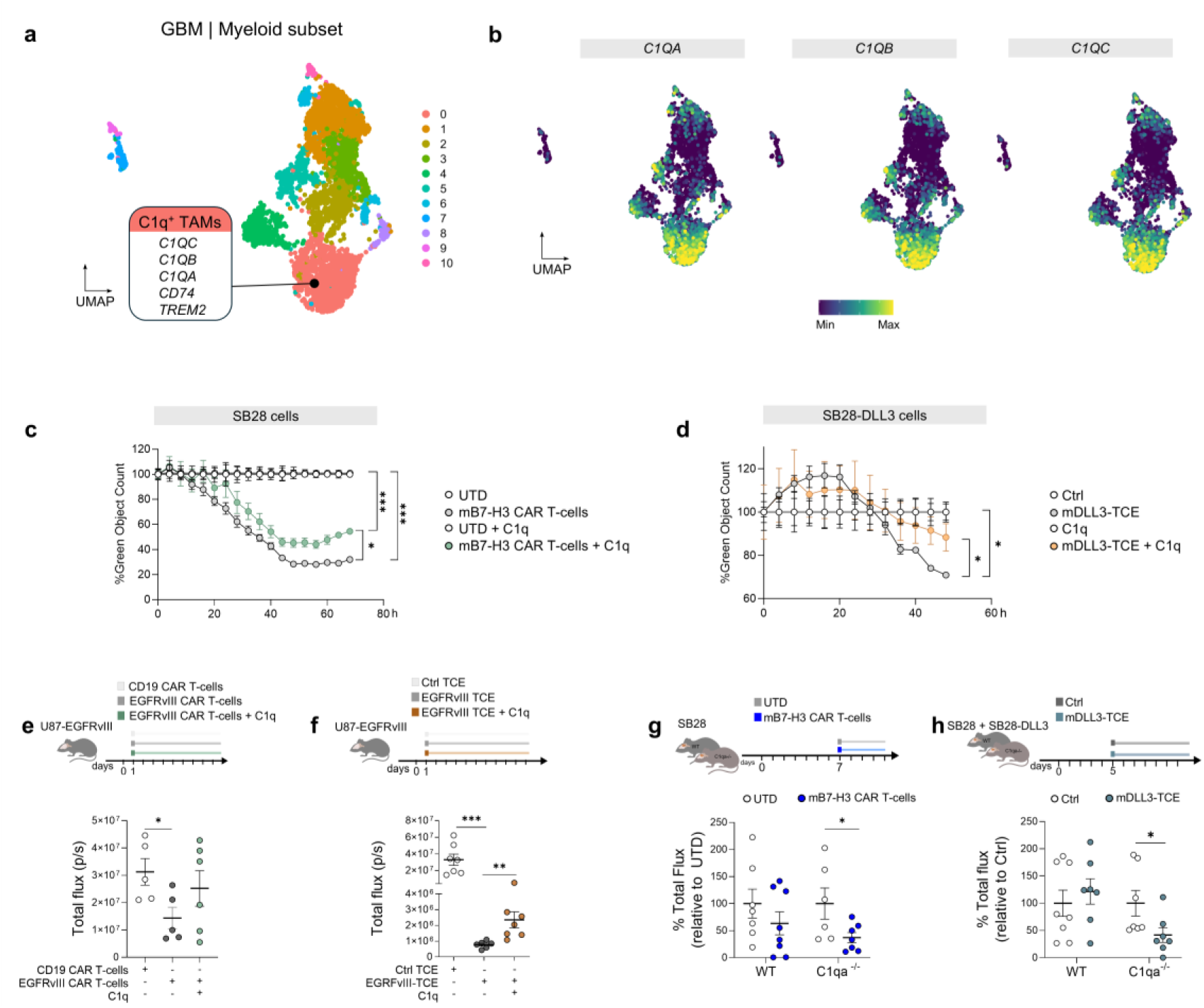
C1q impairs the anti-tumoral response to TCEs and CAR T-cells. **a,** Uniform Manifold Approximation and Projection (UMAP) of scRNAseq data from 3 GBM patients from (Iurlaro et al. co-submitted), indicating the C1q^+^ TAMs cluster. **b,** UMAP coloured according to expression levels of *C1QA*, *C1QB* and *C1QC*. **c,** Cytotoxic assay of SB28 cells co-cultured with murine untransduced (UTD) T-cells or B7H3 CAR T-cells pre-incubated or not overnight with 100 μg/mL C1q at an effector-to-target (E:T) ratio of 2:1. Total green cell count was normalized to the corresponding UTD condition and t=0. **d,** Cytotoxic assay of SB28-DLL3 cells co-cultured with murine CD3^+^ T-cells and mDLL3-TCE at an E:T ratio of 10:1. When indicated, T-cells were pre-incubated overnight with 100 μg/mL C1q. Total green cell count was normalized to the corresponding control condition and t=0. **e,** Mice were intracranially inoculated with U87-EGFRvIII cells. CAR T-cells were pre-incubated overnight with 100 μg/mL C1q or HSA and administered intravenously the day after surgery. CD19 CAR T-cells were used as control. Tumor growth differences were assessed by bioluminescence imaging (n=5 CD19 CAR T-cells, n=5 EGFRvIII CAR T-cells, n=6 EGFRvIII CAR T-cells+C1q). **f,** Mice were intracranially inoculated with U87-EGFRvIII cells. PBMCs were pre-incubated with 100 μg/mL C1q or HSA overnight and administered intravenously the day after surgery together with EGFRvIII-TCE or control TCE. Tumor growth differences were assessed by bioluminescence imaging (n=7 Ctrl TCE, n=8 Ctrl TCE+C1q, n=6 EGFRvIII-TCE, n=7 EGFRvIII-TCE+C1q). **g,** WT or *C1qa*^-/-^ mice were intracranially inoculated with SB28 cells. Mice were randomized at 5 d.p.i. and treated intracranially with B7H3 CAR T-cells or UTD T-cells as control at 7 d.p.i. Tumor growth differences were assessed by bioluminescence imaging (n=7 WT UTD, n=8 WT CAR T-cells, n=6 *C1qa*^-/-^ UTD, n=7 *C1qa*^-/-^ CAR T-cells). **h,** WT or *C1qa*^-/-^ mice were intracranially inoculated with 50% SB28 cells and 50% SB28-DLL3 cells. Mice were randomized at 4 d.p.i. and treated subcutaneously with DLL3-TCE at 5 d.p.i. Tumor growth differences were assessed by bioluminescence imaging (n=8 per group). Data are represented as mean ± SEM. Statistical analyses two-ways ANOVA (a), (b), Mann–Whitney *t*-test (c), (d) or by Student’s *t*-test (e), (f). \**p*<0.05; \*\**p*<0.01; \*\*\**p*<0.001. Dots indicate biological replicates (c), (d), (e), (f).

We hypothesized that C1q^+^ TAMs could mediate resistance to TCTs such as TCEs and CAR T-cells. In order to test this hypothesis, we first performed gain-of-function experiments by co-culturing SB28 GBM cells with T-cells in the presence of C1q to test its impact on CAR T-cell or TCE activity. For this, we leveraged available mouse CAR T-cells targeting murine B7-H3 (mB7-H3) and a mouse TCE targeting DLL3, a well-known target for TCEs^27,28^. Since SB28 cells endogenously express B7-H3 but not DLL3, parental SB28 cells were used for the CAR T-cell experiment, whereas SB28 cells overexpressing DLL3 (SB28-DLL3) were used for the TCE experiment. A C1q dose of 100 μg/mL was used to reflect natural C1q levels in human blood, which range from 50 μg/mL to 250 μg/mL^29^. C1q was able to repress the cytotoxic effect of both CAR T-cells and the TCE, as measured by Incucyte imaging (Fig. 1c and 1d).

We next assessed C1q inhibitory effect *in vivo*, using an orthotopic xenograft mouse model of GBM, in which U87 cells were engineered to express EGFRvIII (U87-EGFRvIII) and inoculated into the brain of NSG mice. EGFRvIII is an EGFR somatic mutation common in GBM (25-30% incidence) that generates a unique extracellular epitope ideal for tumor cell targeting^30^. We then treated these mice with EGFRvIII-targeting CAR T-cells or an EGFRvIII-targeted TCE combined with human leukocytes, with or without C1q. Both CAR T-cells and the TCE inhibited tumor growth and the anti-tumor effect was impaired in the presence of C1q (Fig. 1e and 1f).

We confirmed the inhibitory effect of C1q on TCTs via a loss-of-function approach by using *C1qa*^-/-^ C57BL/6 mice. Wild type (WT) and *C1qa*^-/-^ mice were intracranially inoculated with parental SB28 cells or a mixture of parental and DLL3-overexpressing SB28 cells (SB28-DLL3) to generate orthotopic syngeneic mouse models of GBM and treated with mouse CAR T-cells targeting B7-H3 or with a TCE targeting DLL3, respectively. Interestingly, we observed that, under stringent experimental conditions, CAR T-cells or the TCE showed no significant anti-tumor activity in WT mice, but significantly reduced tumor growth in *C1qa*^-/-^ mice (Fig. 1e and f), highlighting the role of C1q in suppressing CAR T-cells and TCE anti-tumor activity.

Together, these data indicated that C1q inhibits CAR T-cell and TCE activity.

### C1q inhibits T-cell activation

To further understand the effect of C1q^+^ TAMs on TCTs, we assessed the role of C1q on T-cell activation. Culturing peripheral blood mononuclear cells (PBMCs) in the presence of C1q markedly decreased both CD8^+^ and CD4^+^ T-cell activation as measured by surface expression of CD25 and CD69, as well as secretion of granzyme B and IFNγ. Interestingly, C1q-mediated inhibition of T-cells was rescued by the presence of a fragment of a C1q receptor, LAIR2, that sequesters the C1q protein (Fig. 2a, Fig. S2a). Similar results were obtained using the OT-1 system as an antigen-specific model of T-cell activation. Isolated OT-1 CD8^+^ T-cells activated with the SIINFEKL peptide and cultured in the presence of C1q showed decreased levels of 4-1BB, CD25 and CD69, as well as decreased secretion of granzyme B into the media (Fig. 2b). Of note, C1q treatment also reduced phosphorylation of ERK1/2 and LCK in human T-cells, suggesting an impairment of the T-cell receptor (TCR)-associated signaling (Fig. S2b, S2c). This inhibition was corroborated in Jurkat cells, a well-known model to study TCR signaling^31^, where C1q was able to repress phosphorylation of ZAP70 (Fig. S2d).

**Fig 2.**
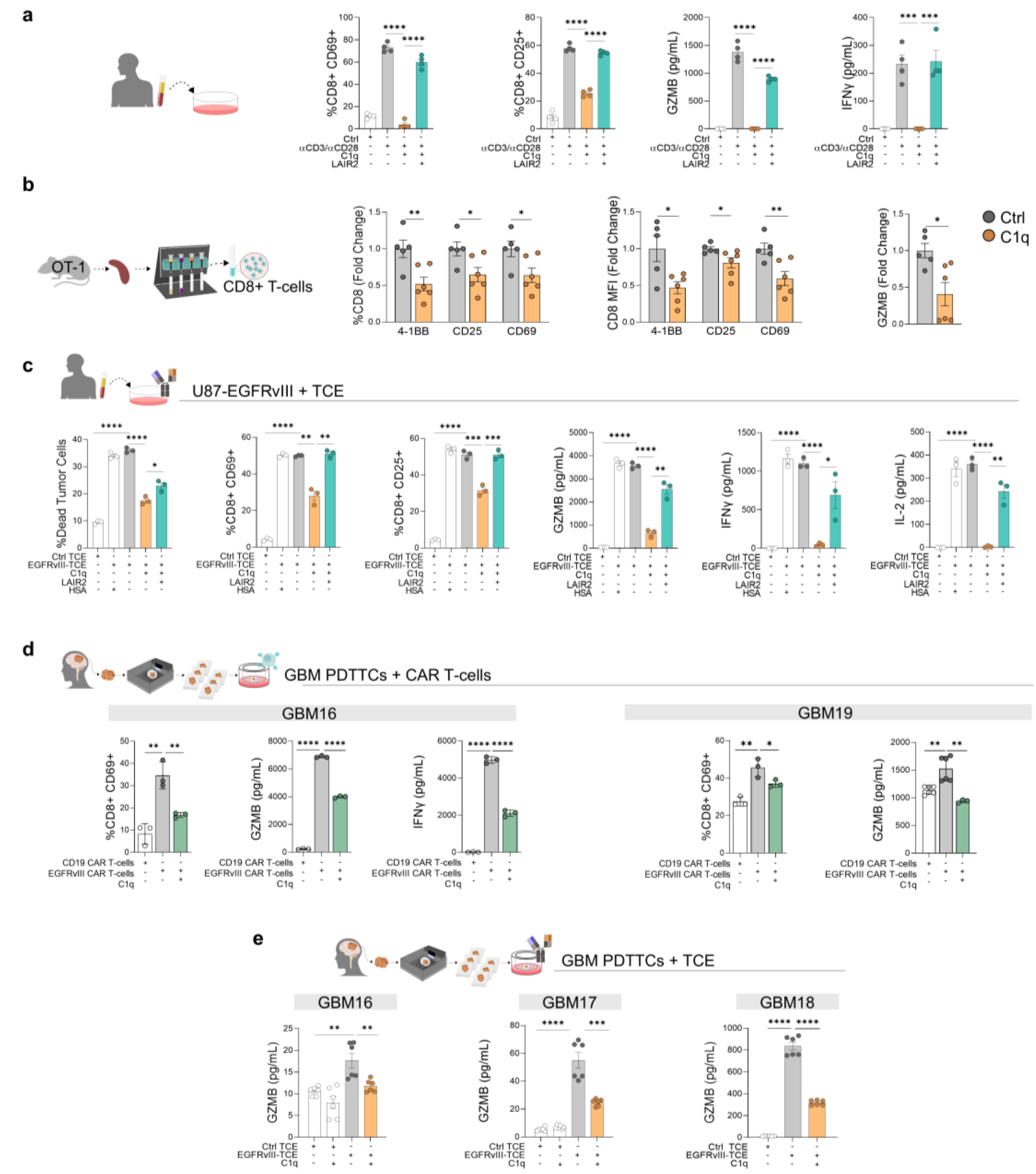
C1q impairs T-cell, TCEs and CAR T-cells activation. **a,** T-cell activation in PBMCs pre-incubated overnight with 100 μg/mL C1q or HSA in the presence or absence of LAIR2 and activated for 24h with soluble αCD3/αCD28. T-cell activation was measured as percentages of CD25^+^ and CD69^+^ cells in CD8^+^ T-cells, as well as ELISA of granzyme B and IFNγ (n=4). **b,** T-cell activation in isolated OT-1 CD8^+^ T-cells pre-incubated overnight with 100 μg/mL C1q and activated for 6h with 1μg/mL SIINFEKL peptide. T-cell activation was measured as percentage and mean fluorescence intensity of the indicated activation markers in CD8^+^ T.cells, as well as ELISA of granzyme B. Data is represented as fold change relative to control (n=5 Ctrl, n =6 C1q). **c,** T-cell activation in PBMCs pre-incubated for 2h with 100 μg/mL C1q or HSA and then co-cultured with U87-EGFRvIII cells. When indicated, C1q and LAIR2 were added to the co-culture prior to the addition of EGFRvIII-TCE or control TCE. T-cell cytotoxicity was measured as tumor cell death by flow cytometry. T-cell activation was measured as percentages of CD69^+^ and CD25^+^ cells in CD8^+^ T-cells, as well as ELISA of granzyme B, IFNγ, and IL-2 (n=3). **d,** T-cell activation in EGFRvIII CAR T-cells pre-incubated overnight with 100 μg/mL C1q or HSA and then co-cultured for 72h with PDTTCs from 2 different GBM patients. CD19 CAR T-cells were used as control. CAR T-cell activation was measured as percentages of CD69^+^ cells in CD8^+^ T-cells, as well as ELISA of granzyme B and IFNγ. **e,** T-cell activation in PBMCs pre-incubated overnight with 100 μg/mL C1q or HSA and then co-cultured for 72h with PDTTCs from 3 different GBM patients together with EGFRvIII-TCE or control TCE. T-cell activation was measured as ELISA of granzyme B. Data are represented as mean ± SEM. Statistical analyses by Student’s t-test (a), (b), (c), (d), (e). \**p*<0.05; \*\**p*<0.01; \*\*\**p*<0.001; \*\*\*\**p*<0.0001. Dots indicate biological replicates (a), (b), (c), or technical replicates (d), (e).

Given our observations that C1q suppresses T-cell activity, we reasoned that a similar phenomenon could happen in T-cells recruited by TCEs. To test this hypothesis, we used the TCE targeting EGFRvIII. U87-EGFRvIII were co-cultured with PBMCs and treated with EGFRvIII-TCE or control TCE. As expected, the EGFRvIII-TCE induced tumor cell death and T-cell activation measured as increase in CD25 and CD69 in CD8^+^ and CD4^+^ T-cells, as well as increased secretion of granzyme B, IFNγ and IL-2, all of which was hampered when PBMCs were pre-incubated with C1q and rescued by the LAIR2 fragment (Fig. 2c, Fig. S2e). Importantly, C1q repressed TCE-dependent phosphorylation of ZAP70, implying that C1q-mediated inhibition of the TCR signaling also occurs in the context of TCEs (Fig. S2f).

To validate our findings in the context of the reality of GBM patients, we used patient-derived tumor tissue cultures (PDTTCs). This model, described in (Iurlaro et al. co-submitted), maintains the architecture of the tissue, as well as the TME including T-cells, myeloid cells, endothelial cells and other cell types, allowing a more accurate study of the regulation of the TME. We co-cultured EGFRvIII-positive GBM PDTTCs (Table S2, S3) with either EGFRvIII CAR T-cells (n=2) or an anti-EGFRvIII TCE (n=3) with or without C1q. Consistent with our previous results, C1q addition decreased CAR T-cell activation as measured by granzyme B, IFNγ and CD69 T-cell-activation marker in both PDTTCs (Fig. 2d) and T-cell activation and cytotoxicity induced by the TCE as measured by the secretion of granzyme B (Fig. 2e).

Taken together, our results show that C1q inhibits T-cell activity, including that induced by TCEs and CARs.

### LIF induces C1q in macrophages

We and others have found that C1q was primarily expressed in myeloid cells in GBM (see Fig. 1a and 1b)^15,19^. We asked why some GBM tumors contain high levels of C1q^+^ TAMs and what are the factors responsible for the presence of this TAM population. Using the GBM cohort from CPTAC^32^, we found LIF to be one of the cytokines that better correlates with C1q^+^ TAMs (Fig. 3a), indicating that LIF could be involved in C1q induction in myeloid cells. To test this, we treated mouse bone marrow-derived macrophages (BMDMs) with recombinant LIF and performed RNAseq. Among other genes, we found that the *C1qa*, *C1qb*, and *C1qc* genes, in addition to other immunosuppressive genes (e.g. *Mrc1* (CD206 protein)), were upregulated by LIF in BMDMs (Fig. 3b). We further validated the RNAseq result by performing a time course experiment analyzing C1q by western blotting (WB) in LIF-treated BMDMs (Fig. 3c). In addition, we demonstrated that the induction of C1q by LIF was dependent on JAK activity, since a JAK inhibitor blunted C1q induction by LIF (Fig. 3d).

**Fig. 3.**
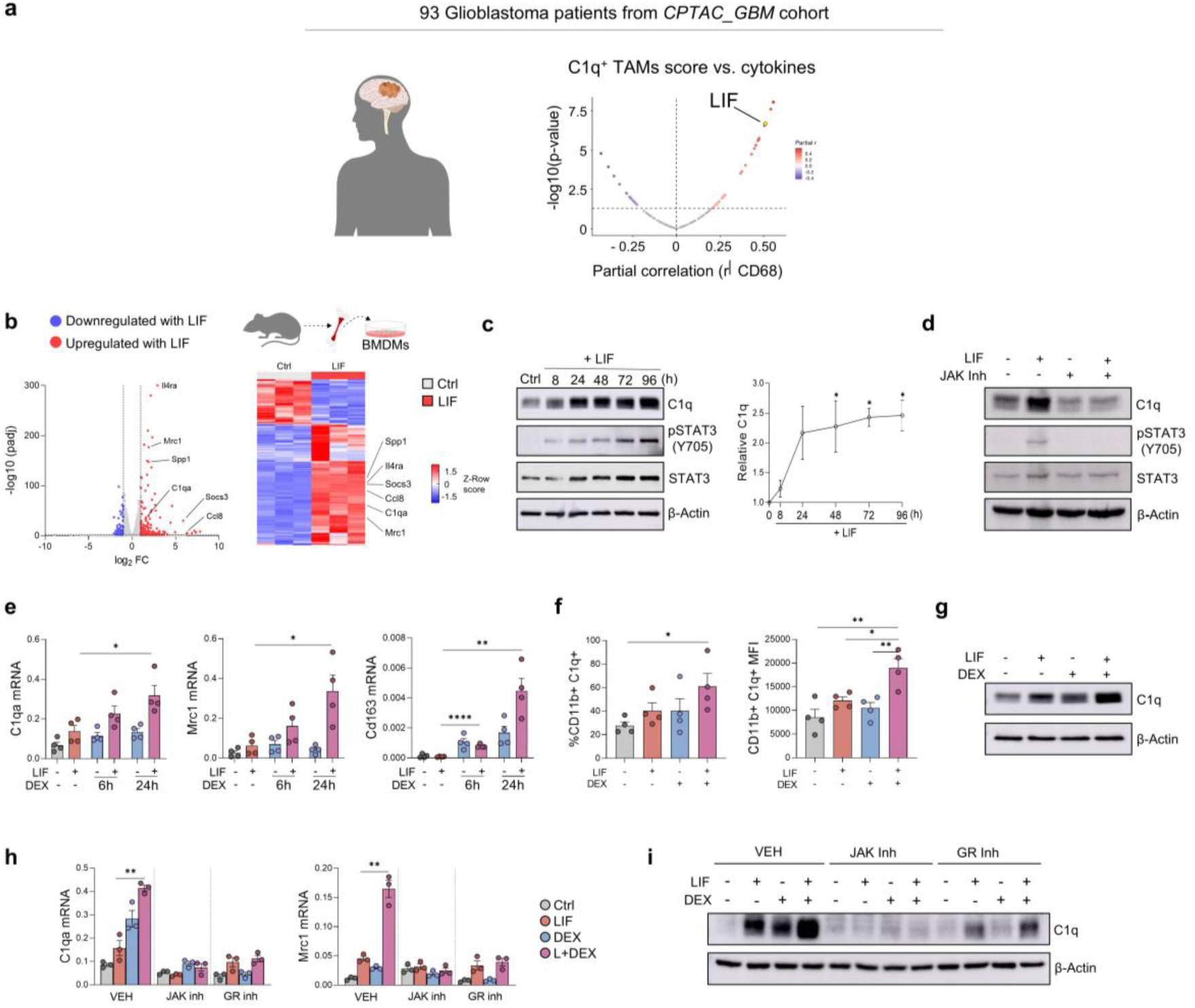
C1q is induced in myeloid cells by a LIF-glucocorticoid axis. **a**, Partial correlation analysis between the C1q⁺ TAM signature score^33^ and cytokine expression controlling for CD68 expression in bulk RNA-seq data from 93 glioblastoma patients (GBM_CPTAC cohort)^32^. **b**, Differential expression analysis of BMDMs treated with 20 ng/mL LIF for 72h (n=3). Left panel, volcano plot of gene expression changes in response to LIF treatment. Upregulated genes in response to LIF are shown in red, while downregulated genes are shown in blue. Right panel, heatmap of the differentially expressed genes in response to LIF with a log2 FC>1 and an adjusted p-value<0.05. **c**, Immunoblot of C1q, pSTAT3 (Y705) and STAT3 in BMDMs control and treated with 20 ng/mL LIF at different time points. β-actin was used as a loading control. Quantification of C1q is represented relative to β-actin and control condition (right panel) (n=3). **d,** Immunoblot of C1q, pSTAT3 (Y705) and STAT3 in BMDMs treated with 20 ng/mL LIF and 0.5 μM JAK inhibitor Tetracyclic Pyridone 6 (P6) for 72h. β-actin was used as a loading control. **e**, mRNA expression of the indicated genes in BMDMs treated with 20 ng/mL LIF for 72h and 100 nM dexamethasone for the last 6h or 24h. Comparison was performed between the LIF and the combination groups (n=4). **f**, Percentage (left panel) and mean fluoresce intensity (right panel) of C1q^+^ cells in BMDMs treated 72h with 20 ng/mL LIF and 100 nM dexamethasone for the last 24h (n=4). **g**, Immunoblot of C1q in BMDMs treated 72h with 20 ng/mL LIF and 100 nM dexamethasone for the last 24h. β-actin was used as a loading control. **h**, mRNA expression of *C1qa* and *Mrc1* in BMDMs treated 72h with 20 ng/mL LIF and 100 nM dexamethasone for the last 24h. When indicated, cells were pre-treated with either vehicle (DMSO), 0.5 μM P6 or 2 μM glucocorticoid receptor inhibitor RU486 prior to the addition of LIF or dexamethasone, respectively. Comparison was performed between the LIF and the combination groups (n=3). **j**, Immunoblot of C1q in BMDMs treated 72h with 20 ng/mL LIF and 100 nM dexamethasone for the last 24h. When indicated, cells were pre-treated with either vehicle (DMSO), 0.5 μM P6 or 2 μM glucocorticoid receptor inhibitor RU486 prior to the addition of LIF or dexamethasone, respectively. β-actin was used as a loading control. Data are mean ± SEM. Statistical analyses by Student’s t-test (c), (e), (f), (h). \**p*<0.05; \*\**p*<0.01; \*\*\*\**p*<0.0001. Dots represent biological replicates (e), (f), (h). Abbreviations: DEX, dexamethasone; VEH, vehicle; GR, glucocorticoid receptor; inh, inhibitor.

Almost all GBM patients are treated with glucocorticoids (usually dexamethasone) at the time of surgical intervention, and glucocorticoids have been described to have an immunosuppressive function^25^. We tested whether dexamethasone impacts the induction of C1q by LIF. Indeed, dexamethasone combination with LIF further increased *C1qa*, *Mrc1* (CD206 protein) and *CD163* expression, at both the transcriptional and protein level (Fig. 3e, 3f, 3g and Fig. S3a). The cooperation of LIF and dexamethasone to induce C1q is dependent on the activity of JAK and glucocorticoid receptor (GR), as JAK and GR inhibitors or GR gene silencing abrogated C1q induction by LIF and dexamethasone (Fig. 3h, 3i and Fig. S3b).

### LIF blockade eliminates C1q^+^ TAMs in human GBM

In order to study the regulation of C1q^+^ TAMs in the context of human GBM, we again took advantage of our PDTTC model. We treated a cohort of PDTTCs from 9 GBM patients (Table S2, Table S3, Fig. 4a) with an anti-LIF neutralizing antibody (mAb) or the isotype IgG control, performed scRNAseq and analyzed the differentially expressed genes (DEGs) in 3 different cell types (malignant cells, lymphoid cells and myeloid cells) for each of the 9 cultures (Fig. S4a). Due to the heterogeneity of GBM, we decided to study each tumor independently and not perform an integrated analysis.

**Fig. 4.**
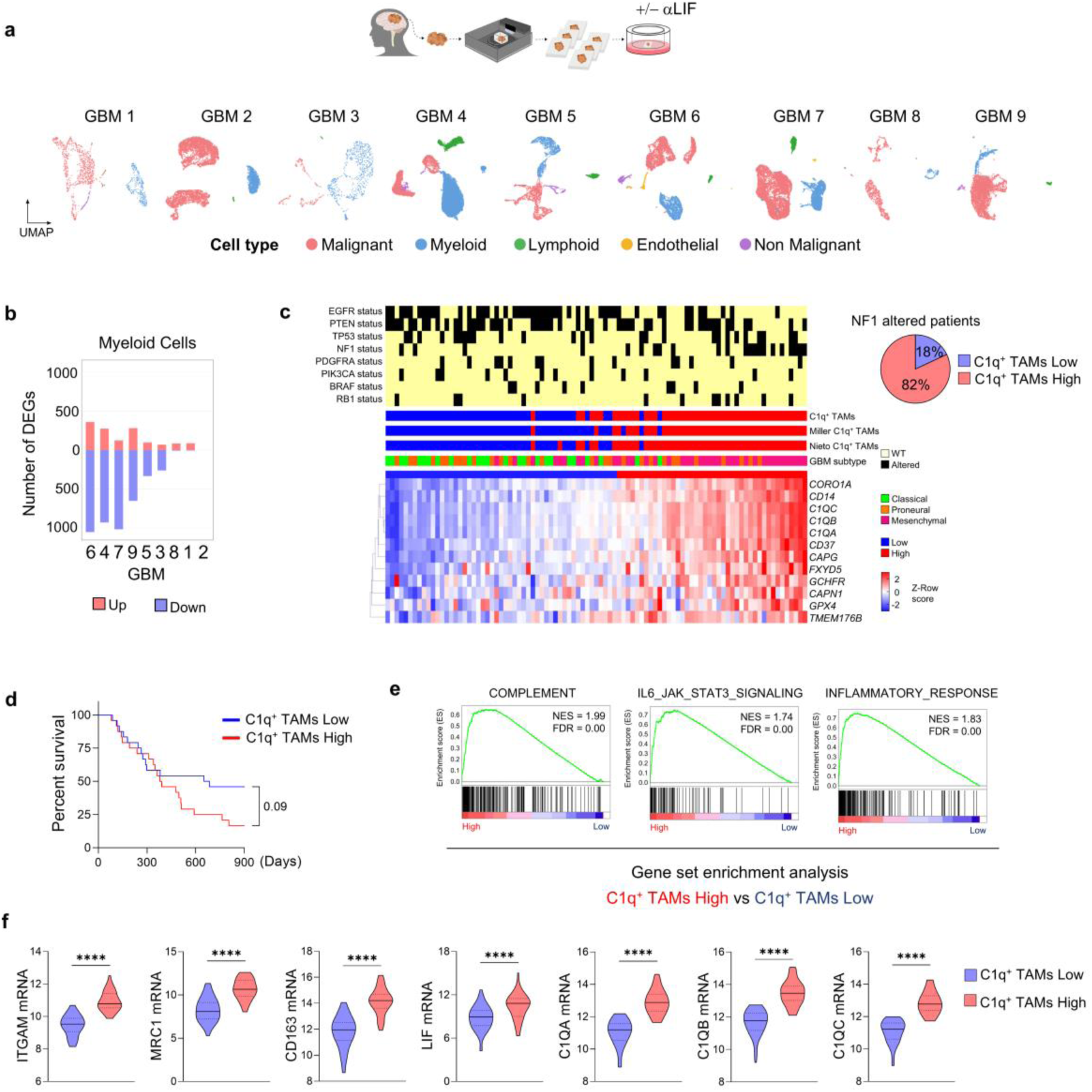
LIF blockade induces profound transcriptional changes in myeloid cells of GBM patients. **a**, Uniform Manifold Approximation and Projection (UMAP) of scRNAseq data from PDTTCs derived from GBM patients (n=9). **b**, Number of DEGs in the myeloid population for each PDTTC sample. **c**, Non-hierarchical k-means clustering analysis of GBM patients from the CPTAC GBM cohort^32^ (n=93) based on the expression of the anti-LIF signature. Heatmap shows the GBM expression subtype of each patient and an overview of significantly altered genes found in at least 5% of samples, the anti-LIF signature expression, and the expression of other C1q^+^ TAMs signatures from the literature^33,34^. Genes with structural variants, fusions, or CNVs were defined as altered. Pie chart of C1q^+^ TAMs signature status in *NF1*-altered patients. **d**, Survival plot of GBM patients from CPTAC GBM cohort stratified by expression levels of the C1q^+^ TAM signature. **e,** Gene Set Enrichment Analysis (GSEA) of indicated gene lists with ranked gene expression list of high vs low C1q^+^ TAM signature groups from the CPTAG GBM Cohort. **f**, Expression of the indicated genes in high and low C1q^+^ TAM signature patients from the CPTAG GBM Cohort (n=93). Statistical analysis by Log rank (Mantel-Cox test) (e), Mann–Whitney t-test (g). ****p<0.0001. Abbreviations: αLIF, anti-LIF

Our first observation was that the transcriptomic response to LIF varied among patients. In accordance with our prior findings that LIF signals through TAMs^22,23^, we found a high number of differentially regulated genes upon anti-LIF treatment in the myeloid compartment, our compartment of interest (Fig. 4b, Fig. S5a). We observed that 4 PDTTCs (GBMs 4, 6, 7 and 9) exhibited a larger number of DEGs in response to anti-LIF, most of them downregulated genes, than the other 5 PDTTCs (Fig. 4b). We identified 74 downregulated gene responses common to GBMs 6, 4 and 7; and 12 gene responses common to GBMs 6, 4, 7 and 9 (Fig. S4a and Table S4). *C1QA*, *B* and *C* were present in both list of genes confirming that anti-LIF downregulated C1q in TAMs. Interestingly, other identified gene responses were previously related to immunosuppressive phenotypes. We considered the 12-gene set as an anti-LIF gene signature.

Interestingly, using GBM tumors from the CPTAC database, we observed that tumors with the C1q^+^ TAM signature overlapped with the ones with the high anti-LIF signature or other published C1q^+^ TAM signatures^33,34^. Moreover, these tumors were enriched for the mesenchymal phenotype and contained *NF1* genomic alterations (Fig. 4c).

In addition, patients with tumors with high C1q^+^ TAM signature exhibited shorter overall survival (Fig. 4d), were generally characterized by having a transcriptome enriched in gene sets related to inflammatory response, complement and IL6-JAK-STAT3 signaling (Fig. 4e) and tended to express high levels of immunosuppressive TAM markers (i.e. *ITGAM*, *MRC1* (CD206 protein) and *CD163*) (Fig. 4f).

We confirmed our results in an independent validation cohort of 6 GBM patients (Table S2). We clustered the 6 GBM tumors based on our C1q^+^ TAM signature and observed that 2 of the GBM tumors (GBMs 14 and 15) clustered together and expressed high levels of the signature (Fig. 5a). Importantly, blockade of LIF decreased the C1q^+^ TAM gene signature in PDTTCs from GBM 15 but not in GBM 12 (Fig. 5b).

**Fig. 5.**
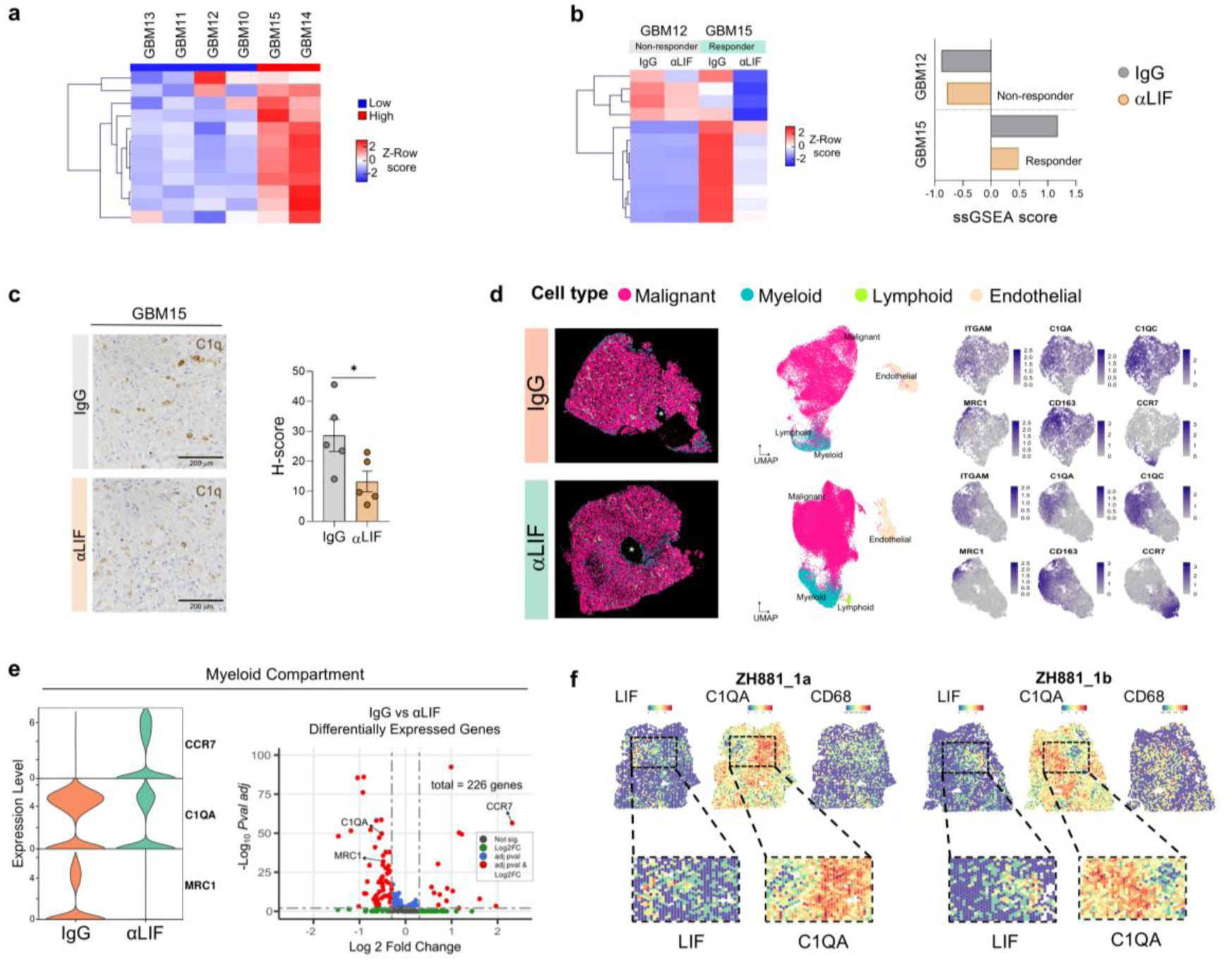
LIF blockade decreases C1q^+^ TAMs. **a**, Non-hierarchical k-means clustering analysis of GBM fresh samples from the validation cohort (n=6) based on the expression of the C1q^+^ TAM signature. **b**, Left panel, C1q^+^ TAM signature transcriptional changes in response to LIF blockade in GBM PDTTCs GBM 12 and GBM 15. Right panel, ssGSEA score for the C1q^+^ TAM signature in both PDTTCs in response to LIF blockade. **c**, Representative IHC of C1q of GBM 15 PDTTCs treated with IgG or anti-LIF. C1q staining was assessed using H-score method. **d**, Single-cell image-based spatial transcriptomics of GBM 15 PDTTCs treated with IgG (top panel) or anti-LIF (bottom panel). The two consecutive slices contain a large vessel marked with an asterisk. Right panel, expression of selected immunosuppressive genes: *C1QA*, *C1QC*, *MRC1* and *CD163*, and the M1-like marker *CCR7* in the myeloid population for both conditions. **e**, Expression levels of the indicated genes in the myeloid population from spatial transcriptomic of IgG and anti-LIF PDTTCs from GBM 15. Right panel, volcano plot representing the differentially expressed genes in the myeloid population. **f**, Spatial maps for samples ZH881_1a and ZH881_1b (10X Visium Spatial Transcriptomics, GSE237183)^35^, highlighting the indicated markers. Bottom panel, zoom in to remark the proximity between the high LIF and high C1q expressing cells. Data are represented as mean ± SEM. Statistical analyses by Student’s t-test (c). *p<0.05.

Consistently, we observed that anti-LIF decreased C1q protein expression in GBM 15 PDTTC by immunohistochemistry (IHC) (Fig. 5c). To further validate the effect of anti-LIF on C1q, we treated two consecutive slices of the PDTTC containing a large vessel with control IgG and anti-LIF and performed spatial transcriptomics using a panel of 300 genes. We focused our studies on TAMs and observed that *C1QA* and CD206 (*MRC1* gene) were decreased upon treatment with anti-LIF, while the M1-like marker *CCR7* was increased (Fig. 5d and 5e).

Taken together, all these results indicated that LIF blockade decreases the levels of C1q and the presence of C1q^+^ TAMs in GBM. Accordingly, reanalysis of publicly available GBM spatial transcriptomics data^35^ showed that in some cases LIF-expressing cells were located at the vicinity of C1q-expressing TAMs, further supporting the relationship between LIF and C1q^+^ TAMs (Fig. 5f).

### A mouse GBM model recapitulates the characteristics of human tumors exhibiting high C1q^+^ TAM and high anti-LIF signatures

We then decided to identify a mouse GBM model that could recapitulate the phenotype of human tumors with high C1q^+^ TAM signature and high anti-LIF signature in order to perform loss-of-function experiments to orthogonally demonstrate the role of C1q in GBM. We developed two syngeneic animal models of GBM by orthotopic inoculation of the mouse GBM cell lines GL261 and its derivative GL261N in immunocompetent C57BL/6 mice. GL261N cells express higher levels of LIF (Fig. 6a) and generated tumors that also expressed high levels of LIF and grew faster than the parental cell line GL261 (Fig. 6a). Notably, the LIF CRISPR/Cas9-knock-out in GL261N generated smaller tumors that grew at a similar rate than GL261 tumors, supporting a pro-tumoral role for LIF in this model (Fig. 6a). Interestingly, GL261N tumors were more infiltrated with CD206^+^ TAMs than GL261 and LIF CRISPR-KO GL261N tumors, similarly as human tumors expressing the anti-LIF gene signature (Fig. 6a). Importantly, GL261N tumors responded to anti-LIF while GL261 tumors did not (Fig. S6a).

**Fig. 6.**
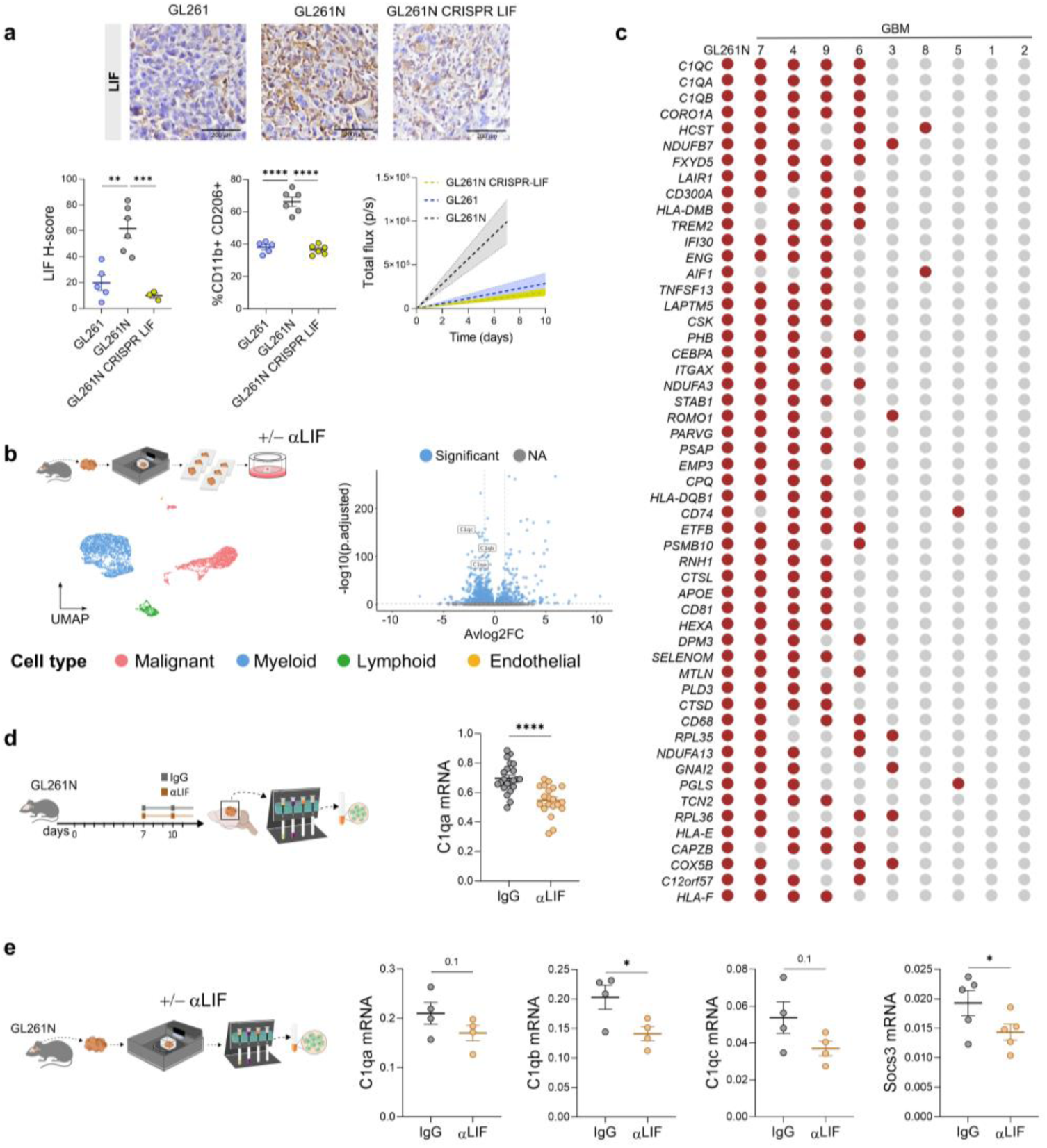
Downregulation of the C1q genes is conserved in murine and human GBM upon LIF blockade. **a**, Representative IHC staining of LIF in GL261, GL261N and GL261N CRISPR LIF tumor sections (top panel). Staining intensity was quantified using the H-score method (bottom left panel). Percentage of CD206^+^ cells in isolated CD11b^+^ cells from each tumor (bottom center panel). Tumor growth assessed by bioluminescence imaging and quantified as total photon flux (p/s) (bottom right panel). (n=5 GL261, n=6 GL261N and GL261N CRISPR LIF). **b**, UMAP of scRNAseq data from MDTTCs derived from a GL261N tumor. Left panel, volcano plot representing the differentially expressed genes (adjusted p-value<0.05) in the myeloid populations in response to anti-LIF blockade. The three C1q genes are marked. **c**, Top shared downregulated genes in response to LIF blockade in the myeloid population between the GL261N MDTTC and the 9 GBM PDTTCs from the discovery cohort. Genes that are downregulated in a particular sample are represented in red. **d**, Mice were inoculated with GL261N, randomized and treated with IgG or anti-LIF from 7 d.p.i. *C1qa* mRNA expression in isolated CD11b^+^ cells from the tumors (n=22 IgG and n=20 anti-LIF). **e**, mRNA expression of the C1q genes in isolated CD11b^+^ cells from IgG or anti-LIF treated MDTTCs from GL261N tumors. *Socs3* included as a control of LIF blockade (n=4). Data are represented as mean ± SEM. Statistical analyses by Student’s t-test (a), (d), (e). \**p*<0.05; \*\**p*<0.01; \*\*\**p*<0.001; \*\*\*\**p*<0.0001. Data are represented as mean ± SEM. Statistical analyses by Student’s t-test (A), Mann–Whitney t-test (B). *p<0.05; **p<0.01; ***p<0.001; ****p<0.0001. Dots replicate biological replicate (a), (d), (e).

We argued that the GL261N tumor model recapitulates the characteristics of human GBM tumors expressing high anti-LIF signature and proceeded to perform mouse-derived tumor tissue cultures (MDTTCs) using the same methodology described for the PDTTCs. Following the same approach used previously, we treated GL261N MDTTCs with anti-LIF or the isotype control IgG and performed scRNAseq and a DEG analysis. We observed a profound change in the transcriptome of myeloid cells in response to anti-LIF, recapitulating the results obtained in human PDTTCs (Fig. 6b).

Notably, we observed a large overlap between the human and mouse gene responses to anti-LIF in TAMs. *C1QA*, *C1QB* and *C1QC* were present among the top conserved genes downregulated by anti-LIF in TAMs from both PDTTCs and MDTTCs (Fig. 6c). This indicated that the C1q response to anti-LIF was conserved between human and mouse demonstrating its physiological relevance. We confirmed that C1q expression was circumscribed to myeloid cells in our mouse model (Fig. S7a) and validated the scRNAseq result by observing a decrease in the expression of C1q in myeloid cells isolated from GL261N tumors from mice (Fig. 6d) and GL261N MDTTCs treated with anti-LIF (Fig. 6e).

### Downregulation of C1q by the blockade of LIF promotes T-cell activation and anti-tumor response

Since we previously described that LIF inhibits the activation of T-cells^22^, we next assessed whether this inhibition of T-cell activation by LIF could be mediated through C1q. We engineered GL261N cells to express the OVA antigen and generated orthotopic syngeneic GL261N-OVA tumors. We performed MDTTCs from these tumors and co-cultured the tissue with CD8^+^ T-cells isolated from OT-1 mice, treating the co-cultures with either anti-LIF or the isotype IgG control. The blockade of LIF induced T-cell activation measured by the induction of CD25 and 4-1BB in CD8^+^ T-cells. Importantly, in *C1qa*^-/-^ mice, basal CD8^+^ T-cell activation was higher, mimicking the effect observed with anti-LIF in wild type MDTTCs. Accordingly, in GL261N-OVA MDTTCs derived from *C1qa*^-/-^ mice, treatment with anti-LIF did not further increase the activation of CD8^+^ T-cells, indicating that LIF-induced repression of T-cell activity was in part mediated by C1q (Fig. 7a).

**Fig. 7.**
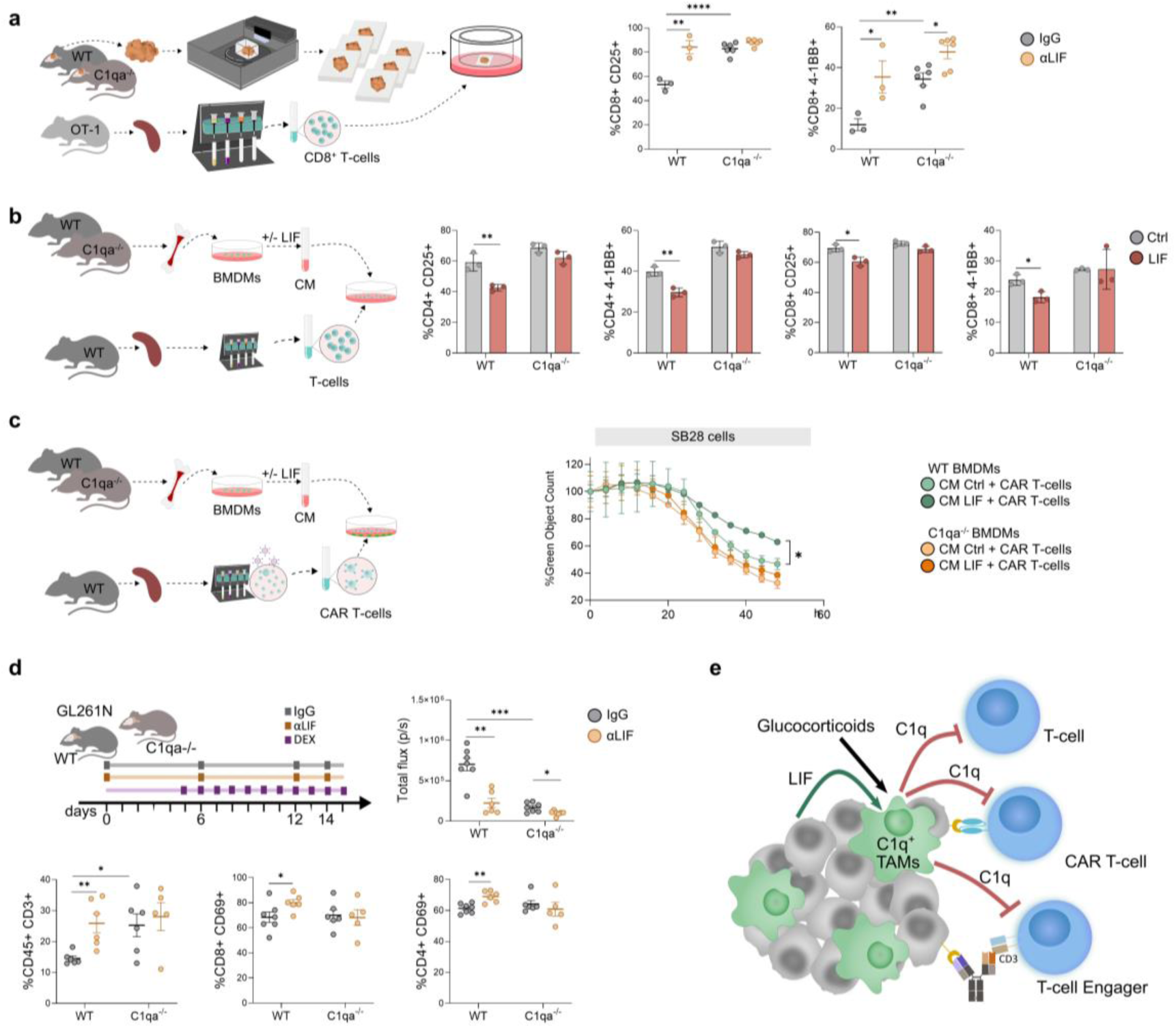
C1q is a mediator of LIF pro-tumoral effect in GBM. **a**, Schematic representation of the experiment (left panel). MDTTCs from GL261N-OVA tumors derived from either WT or *C1qa*^-/-^ mice were co-cultured with CD8^+^ T-cells isolated from the spleens of OT-1 mice and primed with 1 μg/mL of SIINFEKL peptide. T-cell activation in response to anti-LIF was measured as percentage of CD25^+^ and 4-1BB^+^ in CD8^+^ T-cells (n=3 WT, n=6 *C1qa*^-/-^). **b**, Left panel, schematic representation of the experiment. WT or *C1qa*^-/-^ BMDMs untreated or treated with 20 ng/mL LIF were cultured for 72h and conditioned media was collected. T-cells were isolated from the spleen of WT mice, activated with coated αCD3/αCD28 and cultured in the conditioned media (CM) of BMDMs for 24h. T-cell activation was measured as percentages of CD25^+^ and 4-1BB^+^ cells in both CD4^+^ and CD8^+^ T-cells (n = 3). **c**, Schematic representation of the experiment (left panel). Cytotoxic assay of SB28 cells co-cultured with murine B7H3-CAR T-cells at an E:T ratio of 2:1 in the presence of conditioned media from WT or *C1qa*^-/-^ BMDMs control or LIF-treated. Cytotoxicity was assessed by IncuCyte live cell imaging. Total green cell count was normalized to the corresponding UTD condition and t=0. **d**, WT or *C1qa*^-/-^ mice were inoculated with GL261N cells and treated with either IgG or anti-LIF. Dexamethasone was also administered from 5 d.p.i. Tumor growth was assessed by bioluminescence imaging. T-cell infiltration was measured as CD45^+^CD3^+^ cells, and T-cell activation was measured as percentage of CD69^+^ cells in both CD4^+^ and CD8^+^ T-cells (n=7 WT IgG, *C1qa*^-/-^ IgG, *C1qa*^-/-^ anti-LIF, n=6 WT anti-LIF). **e**, LIF and glucocorticoids (e.g. dexamethasone) induce the expression of C1q in TAMs, generating C1q^+^ TAMs. C1q, in turn, inhibits T-cell activation under basal conditions, as well as CAR T-cells and TCE-mediated activation. Data are mean ± SEM. Statistical analyses by Student’s t-test (a), (b), (d). two-ways ANOVA (c), \**p*<0.05; \*\**p*<0.01; \*\*\**p*<0.001; \*\*\*\**p*<0.0001. Dots represent biological replicates (a), (b), (d).

To further study the contribution of macrophage-derived C1q on T-cell activation, we prepared conditioned media from WT and *C1qa*^-/-^ BMDMs treated with LIF to induce the secretion of C1q. As expected, conditioned media from WT BMDMs treated with LIF decreased the activation of T-cells by lowering the levels of CD25 and 4-1BB in CD4^+^ and CD8^+^ T-cells. Importantly, T-cell activation was not affected by conditioned media from *C1qa*^-/-^ BMDMs treated with LIF, confirming that the functional effect of anti-LIF on T-cells is in part mediated by the regulation of C1q expression in macrophages (Fig. 7b).

A similar approach was followed to study the effect of myeloid cell-derived C1q on CAR T-cells activity. B7H3 CAR T-cells were co-cultured with SB28 cells in the presence of conditioned media from control and LIF-treated WT or *C1qa*^-/-^ BMDMs. Notably, conditioned media from LIF-treated WT BMDMs significantly repressed the cytotoxic activity of the CAR T-cells, whereas no effect was observed with conditioned media from LIF-treated *C1qa*^-/-^ BMDMs, supporting our previous findings (Fig. 7c).

Lastly, we generated orthotopic syngeneic GL261N models and treated them with the neutralizing LIF antibody or the isotype IgG control. As we already described, anti-LIF inhibits tumor growth in this model via T-cell mediated cytotoxicity^22,23^. When we performed the same experiment using the *C1qa*^-/-^ mice, we observed that GL261N tumors grew smaller, indicating that C1q exerted a pro-tumoral function in this tumor model. Importantly, anti-LIF had no impact on tumor growth in *C1qa*^-/-^ mice. We dissected the anti-LIF treated tumors and observed that tumors from *C1qa*^-/-^ mice had a higher CD3^+^ T-cells abundance, similar to the infiltration observed in anti-LIF-treated tumors from WT mice. Moreover, anti-LIF did not increase the activation of either CD8^+^ or CD4^+^ T-cells in the context of the *C1qa*^-/-^ (Fig. 7d). Together, these results support that C1q is one of the mediators of anti-LIF anti-tumor response.

## Discussion

GBM is characterized by extreme inter- and intra-tumoral heterogeneity and plasticity, rendering conventional targeted therapies largely ineffective. On the other hand, the immune system has the intrinsic ability to recognize and eliminate abnormal cells regardless of their heterogeneity. Therefore, rationally designed immunotherapies hold the potential to overcome GBM complexity and deliver transformative clinical benefits.

No or minimal therapeutic advances have been achieved in GBM using TCEs and CAR T-cells. One of the primary reasons for this limited success is the immunosuppressive TME in GBM, which impairs the activity of cytotoxic T-cells, whether engaged via TCEs or CARs^9^. This holds true for other solid tumor types besides GBM.

We have discovered that C1q impairs TCE or CAR T-cell activity (Fig. 7e). Our work evidences that C1q expressed in C1q^+^ TAMs acts as a compelling cross-talk factor between the myeloid and adaptive immune systems, with significant implications for cancer immunotherapy. Our findings underscore the importance of the TME in shaping immunotherapy responses and highlight the cause of the differences in TCT treatment response between hematological and solid tumors.

The use of patient-derived models that faithfully recapitulate the GBM TME, PDTTCs, allowed the identification of relevant molecular pathways that could be easily translated into the reality of the patient’s tumors. The immunotherapy field suffers from a lack of robust preclinical models, largely due to limited access to human samples and systems capable of mimicking tumor-immune complexity^14^. By employing a platform that allows systematic and unbiased analysis on a patient-specific basis, we have contributed valuable insights into the determinants of anti-tumor immune responses.

We and others have identified LIF as an important oncogenic factor in GBM^22,23^. Notably, LIF-neutralizing antibodies are currently being tested in clinical trials as anti-cancer agents^36,37^. We identified C1q as an anti-LIF response in TAMs and part of an anti-LIF signature. We observed that the GBMs expressing high C1q^+^ TAM signature also expressed high anti-LIF signature, were predominantly of the mesenchymal subtype and harbored *NF1* mutations. C1q has been implicated as an immunosuppressive factor in cancer, autoimmune diseases, and maternal-fetal tolerance^18^. Interestingly, all three scenarios are also related to LIF function, further supporting the relationship of both factors^24^.

Although C1q⁺ TAMs have been reported in other tumor types and are consistently associated with immunosuppression, the molecular pathways underlying their generation were previously unclear. Clinically, patients with GBM are frequently treated with glucocorticoids (e.g., dexamethasone) to manage inflammation and edema, especially around surgical procedures. While glucocorticoids are known to exert immunosuppressive effects, their mechanisms remain poorly understood. Here, we show that the combined action of LIF and glucocorticoids induces C1q expression and promotes the emergence of C1q⁺ TAMs (Fig. 7e). Nevertheless, further investigation is required to determine whether this mechanism applies to tumors beyond GBM.

In our study, C1q⁺ TAMs emerged as crucial mediators of the immunosuppressive phenotype in GBM impairing the response to CAR T-cells and TCEs. Furthermore, using *C1qa*^-/-^ mice, we demonstrated that C1q is a key mediator of the anti-tumor response to LIF inhibition.

Our findings position C1q and C1q⁺ TAMs as therapeutic targets in isolation or combination with immunotherapies like TCEs and CAR T-cells. These results strongly support the clinical investigation of novel immunotherapeutic strategies targeting the LIF–C1q axis in this devastating disease.

## Methods

### Human Samples

Human glioblastoma (GBM) specimens were obtained on the day of surgical resection from Hospital Universitari Vall d’Hebron and Hospital Clínic (Barcelona, Spain). The clinical protocol was approved by the respective Institutional Review Boards of both centers (Comitè d’Ètica d’Investigació Clínica, CEIC), and written informed consent was obtained from all patients prior to sample collection and use for research purposes.

### Patient- and mouse- derived tumor tissue cultures

For the generation of PDTTCs, tumor samples were first processed into smaller pieces, which were then embedded in agarose (Promega). Agarose-embedded tumor pieces were cut into 350 μm thick sections using a vibratome (Leica VT 1200S), and slices were then transferred into 0.4 μm cell culture inserts (Millipore) within 6-well plates or 24-well plates containing 1.2 mL or 0.6 mL, respectively, of Neurobasal medium (Life Technologies) supplemented with B-27 (Life Technologies), penicillin/streptomycin (Life Technologies) and growth factors (20 ng/mL EGF and 20 ng/mL FGF-2) (PeproTech). Human anti-mouse/human LIF blocking antibody (referred to as anti-LIF, developed in house) or its corresponding control human IgG (BioXCell) was added to the culture on day 1 at a concentration of 10 µg/mL. The cultures were kept at 37°C with constant humidity, 95% air and 5% CO_2_ for 48 to 96 hours.

For the generation of MDTTCs, the same procedure was followed. Briefly, tumors were resected from mice, embedded in agarose and sectioned in 350 μm thick slices, which were then cultured with either 10 µg/mL anti-LIF or IgG for 48 to 72 hours.

PDTTCs or MDTTCs were then enzymatically digested using the Human Tumor Dissociation kit or Mouse Tumor Dissociation kit (Miltenyi Biotec), respectively, prior to proceeding with flow cytometry analysis or scRNAseq. Media was collected for ELISA. In the case of FFPE generation, tumor slices were fixed in 4% paraformaldehyde (WVR) solution overnight and then transferred to a 70% ethanol (Vidrafoc) solution for paraffin embedding.

### Primary cell cultures

Bone marrow derived macrophages (BMDMs) were obtained from 6 to 10-week-old female or male WT or *C1qa*^-/-^ C57BL/6 mice as previously described^22^. Briefly, bone marrow cells isolated from the humerus, femur and tibia were cultured in DMEM medium (Life Technologies) supplemented with 20% heat-inactivated FBS (Life Technologies) and L-cell conditioned medium as a source of macrophage-colony stimulating factor. For the generation of L-cell conditioned medium, L929 cells were cultured in DMEM medium supplemented with 10% heat-inactivated FBS for 7 days, after which supernatant was collected and stored in aliquots at -20°C until use. After 6 days of culture, differentiated macrophages were collected and seeded for subsequent experiments in DMEM medium supplemented with 1% penicilin/streptomycin.

For the culture of CD8^+^ T-cells, single-cell suspensions were prepared from the spleens of OT-1 mice by mechanical disruption, and cells were isolated using the CD8a^+^ T Cell Isolation Kit and the MultiMACS Cell24 separator Plus (Miltenyi Biotec). Isolated CD8^+^ T-cells were then cultured in RPMI medium supplemented with 10% heat-inactivated FBS, 1% penicillin/streptomycin, 10 mM HEPES (Sigma Aldrich), 1 mM Sodium Pyruvate (Life Technologies), 0.1 mM Non-Essential Aminoacids (Life Technologies), 50 μM β-mercaptoethanol (Sigma) and mIL-2 (R&D Systems). For the culture of CD3^+^ T-cells, the same protocol was followed using spleens from WT C57BL/6 mice and the Pan T Cell Isolation Kit II (Miltenyi Biotec). For the conditioned media experiment, isolated CD3^+^ T-cells were cultured in conditioned media of WT or *C1qa*^-/-^ BMDMs and mIL-2 (R&D Systems) and activated with 2 μg/mL coated αCD3 and αCD28 (Ultra-LEAF^™^ Purified anti-mouse CD3ε Antibody and Ultra-LEAF^™^ Purified anti-mouse CD28 Antibody, Biolegend).

Peripheral blood mononuclear cells (PBMCs) were obtained from whole blood by centrifuge density separation using Lymphosep (Biowest). CD3^+^ T-cells were isolated from healthy donors’ buffy coats using RosetteSep Kits (Stem Cell Technologies). PBMCs and isolated CD3^+^ T-cells PBMCs were cultured in RPMI media supplemented with 10% heat-inactivated FBS, 1% penicillin/streptomycin and hIL-2 (R&D Systems).

### Cell lines

GL261 cells were obtained from DMSZ (ACC 802). GL261 cells were infected with a lentiviral construct expressing luciferase (pLenti CMV Luciferase; Addgene) to allow the detection of cells *in vivo*. GL261N cells were derived from GL261^22^. For the generation of GL261N CRISPR/LIF cells, the commercial mouse LIF CRISPR/Cas9 KO plasmid (Santa Cruz Biotechnology) was used as previously described^22^. For the generation of GL261N-OVA cells, a pRRL-OVA plasmid kindly provided by Dr. Recio Conde was used. First, HEK293T cells were transfected using Lipofectamine 2000 (Invitrogen) with pMD2.G enveloping plasmid, psPAX.2 packaging plasmid and the OVA plasmid. The following day, media was replaced, and supernatants containing the lentivirus were collected after 24 hours. GL261N cells were then infected with medium containing lentivirus and polybrene (Sigma) at a concentration of 8 µg/mL. After 16 hours, cells were washed, and medium was replaced. Infected cells were selected using Blasticidin S HCl solution (Santa Cruz). All cell lines were cultured in RPMI medium supplemented with 10% FBS and 1% penicillin/streptomycin.

U87 cells were purchased from ATCC (HTB-14). U87-EGFRvIII cells were generated by stably transfecting the U87 cells with human EGFRvIII^30^. Both cell lines were cultured in DMEM media supplemented with 10% FBS and 1% penicillin/streptomycin.

Jurkat cells were kindly gifted by Dr. Sonia Guedan and cultured in RPMI media supplemented with 10% heat-inactivated FBS, 1% penicillin/streptomycin and 1% HEPES.

SB28-Ohlfest cells expressing GFP and luciferase were obtained from DMSZ (ACC 880). For the generation of SB28-DLL3, murine DLL3 was cloned into a lentiviral vector (VectorBuilder), and lentivirus were produced in HEK293T cells as previously described. SB28 cells were then infected with medium containing lentivirus and polybrene (Sigma) at a concentration of 8 µg/mL. Cells overexpressing DLL3 were selected by sorting (Cytek^®^ Aurora CS spectral cell sorter). All cell lines were cultured in DMEM media supplemented with 10% FBS and 1% penicillin/streptomycin.

### Treatments

The following treatments were used as indicated for each experiment: murine recombinant LIF (Millipore), Dexamethasone (Sigma Aldrich), Tetracyclic pyridone 6 (P6, Sigma-Aldrich), RU486 (MedChem), native human C1q protein (Abcam), Recombinant Human LAIR2-Fc Chimera (BioLegend), Human Serum Albumin (HSA, Sigma-Aldrich), EGFRvIII-TCE/ control TCE, mDLL3-TCE.

### siRNA transfection

BMDMs were seeded and treated when indicated with 20 ng/mL LIF. After 24 hours, cells were transfected with 30 nM glucocorticoid receptor siRNA (ON-TARGETplus siRNA Nr3c1 Mouse, Dharmacon) or non-targeting siRNAs as control (ON-TARGETplus Non-targeting Control siRNAs, Dharmacon) using Lipofectamine 3000 Transfection Reagents, following the manufacturer’s protocol (Life Technologies). LIF treatment was maintained during transfection. The following day, cells were washed with PBS and fresh medium was added. BMDMs were then treated with 100 nM dexamethasone. LIF treatment was also renewed. Cells were collected the following day for subsequent analysis. Silencing efficacy was measured by qPCR.

### CAR T-cell generation

Human CD19 CAR T-cells were a kind gift from Sonia Guedan’s group^38^. For the generation of human EGFRvIII CAR T-cells, T-cells were first isolated from healthy donors using RosetteSep Kits (Stem Cell Technologies) and pre-activated with human CD3/CD28 Dynabeads (ThermoFisher) at a 2:1 bead to T-cell ratio in non-treated 6-well plate at a concentration of 1×10^6^ cells/mL. The following day, T-cells were infected with a lentiviral pCDCAR1 plasmid containing EGFRvIII 3C10 scFv (3C10), CD8α hinge, 4-1BB costimulatory, and CD3z signaling domain (Creative Biolabs). After 48 hours of lentiviral transduction, dynabeads were removed and CAR T-cells were expanded in the presence of 10 ng/mL human IL-7 and IL-15 (Miltenyi Biotec).

Murine B7H3 CAR construct was cloned into a retroviral pMSCV plasmid containing B7H3 m8524 scFv, mCD8α hinge and TM, mCD28 co-stimulatory and mCD3z signaling domain (VectorBuilder). Retrovirus were produced by transfecting 12.5×10^6^ PhoenixEco cells (ATCC; CRL-3214) with 22.5 μg of the CAR retroviral vector and 10.5 μg of pCL-Eco (kindly gifted by Dr. Marcela Maus) with 150 μL of Lipofectamine 2000. For the generation of murine B7H3 CAR T-cells, T cells were isolated from mechanically disrupted spleens of WT C57BL/6 mice using EasySep™ Mouse T Cell Isolation Kit (Stem Cell Technologies). T-cells were then activated with murine CD3/CD28 Dynabeads (Thermofisher) at a 1:1 bead to T-cell ratio in mouse T-cell media supplemented with 50 U/mL human IL-2 and 10 ng/mL human IL-7 for 48 hours. After bead removal, T-cells were placed in the presence of 10 ng/mL human IL-7 and IL-15 and incubated with CAR retrovirus or empty retrovirus in fibronectin-coated plates (Takara Bio) for 72 hours.

CAR expression was assessed by flow cytometry with biotin-anti-human IgG (Jackson ImmunoResearch) plus APC-streptavidin (BD Bioscences).

### Co-culture experiments

For the EGFRvIII-TCE co-culture with PDTTCs or U87-EGFRvIII-derived MDTTCs, tumor slices were co-cultured in the presence of 100 nM dexamethasone with 1×10^7^ PBMCs and 10 µg/mL control TCE or EGFRvIII-TCE for 72 hours^30^. For EGFRvIII CAR T-cells experiments, PDTTCs were co-cultured with 3×10^6^ CD19 CAR T-cells or EGFRvIII CAR T-cells for 72 hours. When indicated, PBMCs or CAR T-cells were pre-incubated overnight on C1q or HSA (100 µg/mL) coated plates before proceeding with the co-culture. The corresponding PBMCs plus TCEs or CAR T-cells were added directly on top of the tumor slices together with C1q.

For U87-EGFRvIII cells co-culture experiments, 1×10^6^ PBMCs were co-cultured with 1×10^5^ U87-EGFRvIII cells in a 10:1-effector-to-target (E:T) ratio in RPMI medium supplemented with 10% heat-inactivated FBS, in the presence of 10 ng/mL control TCE or EGFRvIII-TCE. When indicated, PBMCs were pre-incubated for 2 hours on C1q or HSA (100 µg/mL) coated plates prior to co-culture. For the detection of pZAP70, CD3^+^ T-cells were incubated overnight with 100 μg/mL human C1q or HSA and co-cultured for 1 hour with U87-EGFRvIII cells in a ratio of 1:1 E:T (2×10^5^ CD3^+^ T-cells:2×10^5^ U87-EGFRvIII) in the presence of 1 μg/mL EGFRvIII-TCE or control TCE.

For the GL261N-OVA MDTTCs co-culture experiments, MDTTCs were treated with 10 µg/mL anti-LIF or IgG and 100 nM Dexamethasone 24 hours prior to the co-culture. In parallel, isolated CD8^+^ T-cells from OT-1 mice were primed overnight with 1 μg/mL SIINFEKL peptide (Invivogen) and mIL-2 (R&D Systems). Cells were then withdrawn from activation and maintained in culture until being added on top of the tumor slices for 48 hours.

### Cytotoxicity assays

For cytotoxicity assays, 2.5×10^5^ GFP⁺ SB28 or SB28-DLL3 target cells were seeded in flat-bottom 96-well plates. SB28 cells were co-cultured with murine B7H3 CAR T-cells or UTD T-cells at an E:T ratio of 2:1. SB28-DLL3 cells were co-cultured with murine CD3⁺ T-cells at an E:T ratio of 10:1 in the presence or absence of 10 μg/mL murine DLL3-TCE. When indicated, T-cells or CAR T-cells were pre-incubated overnight with 100 µg/mL C1q.

For co-culture experiments with BMDM conditioned media, SB28 cells were co-cultured with murine B7H3 CAR T-cells or UTD T-cells in 50% conditioned medium derived from control or LIF-treated WT or *C1qa*^-/-^ BMDMs, and 50% mouse T-cell medium.

Plates were incubated in an IncuCyte SX5 live-cell analysis system (Sartorius). Images were acquired every 4 hours using a 10× objective, capturing four fields per well. Cytotoxicity was quantified using IncuCyte SX5 analysis software based on GFP signal.

### Animal studies and *in vivo* therapies

All animal experiments were approved by and performed according to the guidelines of the Institutional Animal Care Committee of the Vall d’Hebron Research Institute in agreement with the European Union and national directives under the CEEA 20/23 and 85/17. C57BL/6 mice (C57BL/6JRj) were purchased from Janvier. *C1qa*^-/-^ mice were initially purchased from The Jackson Laboratory (Strain #:031675) and bred in our animal facility to establish a colony. C57BL/6-Tg (TcraTcrb)1100Mjb/J (C57BL/6 background, referred as to OT-1) mice were kindly gifted by Dr. Francisco Barriga. NOD.Cg-Prkdcscid Il2rgtm1Wjl/SzJ (NOD scid gamma, NSG) mice were purchased from Charles River.

For syngeneic brain tumor models, 3×10^5^ GL261, GL261N, GL261N CRISPR/LIF or GL261N-OVA cells, 1×10^4^ SB28 cells or a mix of 5×10^4^ SB28 and 5×10^4^ SB28-DLL3 cells, all of them expressing luciferase, were stereotactically inoculated into the corpus striatum of the right brain hemisphere (1 mm anterior and 1.8 mm lateral to the lambda; 2.5 mm intraparenchymal) of the mice. When indicated, mice were randomized at 7 days post inoculation (d.p.i) into different treatment groups and 300 µg anti-LIF or control IgG was administered intraperitoneally twice a week. Alternatively, mice were administered intraperitoneally with 300 µg anti-LIF or control IgG twice a week from the day of the surgery. Dexamethasone (Fortecortin, ERN) was administered daily intraperitoneally from 5 d.p.i at a dose of 2 mg/kg/day when indicated. For the experiments with murine CAR T-cells, 1×10^6^ B7H3 CAR T-cells or UTD T-cells were intracranially inoculated at 7 d.p.i. For the experiments with murine DLL3-TCE, 1 mg/kg DLL3-TCE was administered subcutaneously at 5 d.p.i. Tumor progression was monitored by bioluminescence measurements using the Xenogen IVIS^®^ Spectrum. Mice were euthanized when they exhibited clinical signs of disease or distress and tumors were collected and used for MDTTCs or for further analysis. For the isolation of CD11b^+^ cells, tumors were enzymatically digested with Mouse Tumor Dissociation kit, and cells were isolated using CD11b MicroBeads and the MultiMACS Cell24 separator Plus (Miltenyi Biotec).

For the experiments with EGFRvIII-TCE and EGFRvIII CAR T-cells, 1.5×10^5^ U87-EGFRvIII cells expressing luciferase were intracranially inoculated in 5-week-old female NSG mice. After 1 day, 1×10^7^ PBMCs together with 5 mg/kg of EGFRvIII-TCE or Control TCE were i.v. inoculated into the mice and treatment continued twice a week subcutaneously. For CAR T-cell experiments, 3.5×10^6^ CD19 or EGFRvIII CAR T-cells were i.v. inoculated into the mice at 1 d.p.i. When indicated, PBMCs or CAR T-cells were pre-incubated overnight on C1q or HSA (100 µg/mL) coated plates before proceeding with the *in vivo* injection. Tumor progression was monitored by bioluminescence measurements using the Xenogen IVIS^®^ Spectrum.

### Flow Cytometry

For murine experiments, digested tumors or MDTTCs were strained through a 50 μm filter (Sysmex) and resuspended in PBS. Fc-block was performed with anti-CD16/32 (BD Pharmingen) for 20 minutes. In the case of isolated CD11b^+^, BMDMs and isolated T-cells, the blocking step was not performed. Staining for cell surface markers was performed for 15 minutes using antibodies against CD45 (BioLegend; 1:222), CD3 (BioLegend; 1:100), CD8 (BD; 1:40), CD4 (BD; 1:250), CD11b (BD 1:100), F4/80 (BD 1:25), LY6G (BD; 1:33), CD25 (Invitrogen, BD; 1:100), CD69 (BD; 1:100), CD206 (BD; 1:80) and 4-1BB (BioLegend; 1:100).

For intracellular staining, anti-C1q (Abcam; 1:100) was used together with the Intracellular Fixation & Permeabilization Buffer Set (eBioscience) according to manufacturer’s instructions, followed by staining with secondary donkey anti-rabbit FITC-IgG antibody (Invitrogen).

For human experiments, digested PDTTCs or MDTTCs from co-cultures experiments with human PBMCs were strained through a 50 μm filter (Sysmex) and resuspended in PBS. Non-specific Fc Receptor-mediated antibody binding was blocked using Human BD Fc Block (BD Pharmigen) for 20 minutes. In the case of isolated PBMCs, the blocking step was not performed. Staining for cell surface markers was performed for 15 minutes using antibodies against CD45 (BD; 1:100), CD3 (BD; 1:100), CD8 (BD; 1:100), CD4 (BioLegend; 1:100), CD25 (BD; 1:100) and CD69 (BD; 1:100).

In all cases, samples were previously incubated for 20 minutes with Fixable Viability Stain 620 (BD; 1:2000) or LIVE/DEAD™ Fixable Yellow Dead Cell Stain Kit (ThermoFisher; 1:2000) to assess viability. Gating was performed inside viable cells unless otherwise indicated.

For pZAP70 staining, the Phosflow Fix and Perm/Wash buffers (BD) were used according to manufacturer instructions. In the case of Jurkat cells, following overnight incubation with 100 μg/mL C1q or HSA, cells were washed, incubated with 1 μg αCD3 and 1 μg αCD28 for 20 minutes at 4°C, washed again and incubated 3 minutes at 37°C before fixation. Staining was performed for 30 minutes in the dark using an antibody against pZAP70 (BD; 1:5).

Samples were acquired on a BD FACSCelesta^™^ flow cytometer and data analyzed with Flow Jo software (v10.8.1).

### Immunoblot

Cells were lysed in ice-cold RIPA buffer (50 mM Tris-HCl pH 7.4, 150 mM NaCl, 1% NP40, 0.5% Deoxycolate, 0.1% SDS, 10 Mm NaF, 20 mM β-Glycerolphosphate) supplemented with phosphatase inhibitors cocktails number 2 (Sigma Aldrich) and number 3 (Sigma Aldrich). Protein lysates were sonicated and cleared by centrifugation and concentration was determined using the BCA Protein Assay Kit (ThermoFisher). Proteins were resolved in SDS-PAGE gels using the Mini Protean III system (BioRad), and electrotransferred onto nitrocellulose membranes (iBlot^™^ 3 Transfer Stacks Mini NC, Invitrogen) using a semi-wet blotting system (iBlot^™^ 3 Western Blot Transfer System, Invitrogen). Following incubation with the corresponding primary antibody and secondary antibodies, detection was performed using enhanced chemiluminescence (ECL) reagent Immobilon^®^ Western Chemiluminescence HRP Substrate, a luminol-based HRP substrate, and captured in the iBright Imaging System (Invitrogen, ThermoFisher Scientific). The primary antibodies used for immunoblotting were as follows: anti-C1q (Abcam), anti-p-STAT3 (CellSignaling), anti-STAT3 (CellSignaling), anti-p-LCK (Sigma Aldrich), anti-LCK (CellSignaling), anti-β-actin (SIGMA), anti-p-ERK1/2 (CellSignaling), anti-ERK1/2 (CellSignaling) and anti-GAPDH (R&D Systems). The secondary antibodies used were as follows: anti-Rabbit IgG HRP Linked (Cytiva), anti-Mouse IgG HRP Linked (Cytiva).

### ELISA

To quantitative determine the proteins levels of mouse LIF, mouse Granzyme B, human Granzyme B, human IFNγ and human IL-2 secreted to the media, Mouse or Human Duo-Set ELISA kit (R&D systems) were used following manufacturer’s specifications.

### Quantitative real-time PCR

RNA was extracted from cells or previously disgregated tumor samples using the RNeasy Mini or Micro kit (Qiagen). Reverse transcription was performed for cDNA formation using iScript Reverse Supermix (BioRad) according to the manufacturer’s instructions. RT-qPCR reaction was performed using Taqman probes (Applied Biosystems) according to the manufacturer’s instructions and carried out in a CFX384 Touch^™^ Real-Time PCR Detection System (BioRad). Results were expressed as fold change calculated using the ΔΔCt method relative to the control sample. Murine or human ACTB were used as internal normalization controls.

### Immunohistochemical and immunofluorescence staining

For immunohistochemical staining, formalin-fixed Paraffin-Embedded (FFPE) tissue sections were deparaffinized overnight at 60°C, followed by rehydration through xylene (PanReac Applichem) and graded ethanol (Vidrafoc). Antigen retrieval was performed using pH 6 or pH 9 Citrate solution (DAKO), followed by microwave heating for 20 minutes. Sections were then incubated for 10 minutes with 10% peroxidase (Sigma) and blocked with 3% BSA in 0.01% Tween-TBS for 1 hour at room temperature. Slides were incubated overnight at 4°C in a humid chamber with the corresponding primary antibodies: rabbit anti-C1q antibody (antigen retrieval pH 9, Abcam; 1:500); mouse anti-LIF antibody (antigen retrieval pH 6, Atlas Antibodies; 1:300). Detection was performed using EnVision system for mouse antibodies (DAKO) or the rabbit-specific IHC kit (Abcam), following manufacturer’s instructions. Hematoxylin (Merck) was used for counterstaining, followed by dehydration through graded ethanol and xylene. Slides were mounted with DPX (Sigma-Aldrich) and air dried prior to image acquisition using a Nikon Eclipse Ci microscope. Analysis was performed using QuPath and quantified by the H-score formula: 3×% high + 2×% intermediate + 1×% low staining intensity.

For co-immunofluorescence staining, FFPE tissue slides were deparaffinized overnight at 60°C and rehydrated as previously described. Antigen retrieval was performed using pH 6 Citrate solution. Permeabilization was performed by incubating slides in 2% Tween-PBS for 30 minutes at room temperature. Blocking was performed in 3% BSA diluted in Tween-PBS + 10% Donkey Serum for 1 hour at room temperature. Slides were incubated overnight at 4°C with the corresponding primary antibodies diluted in PBS 3% BSA: rabbit anti-CD11b (Abcam; 1:500), mouse anti-C1q (Abcam; 1:100). The following day, slides were incubated with the appropriate secondary antibodies diluted 1:200 in PBS for 2 hours at 4°C: anti-rabbit IgG AlexaFluor488 conjugated (Invitrogen) and donkey anti-mouse IgG AlexaFluor568 conjugated (Invitrogen). Slides were mounted in ProLong Glass antifade mountant with NucBlue stain (ThermoFisher), and images were taken in a confocal microscope Zeiss LSM980 and processed using ImageJ software.

### Statistical analysis

All results are represented as mentioned in figure-legends, where the mean ± standard deviation (SD) or standard error of the mean (SEM) is shown. Data were analyzed using GraphPad Prism 8.0 software. For comparisons among two different groups, the p-value (*p*) using Student’s *t*-test (paired or unpaired) for parametric variables or Mann–Whitney test for non-parametric variables was calculated. For multiple independent groups, two-way ANOVA was performed as a parametric test. For survival tests, Mantel-Cox statistics were used. Significant outliers were identified using GraphPad Prism and excluded from the subsequent analysis. For statistical significance. \**p*<0.05, \*\**p*<0.01, \*\*\**p*<0.001, \*\*\*\**p*<0.0001, n.s. not significant.

### Single cell RNAseq data analysis

#### Preprocessing

FASTQ files derived from sequencing were processed by 10X Genomics software Cell Ranger using a container (https://hub.docker.com/r/litd/docker-cellranger) to quantify and align the reads to Homo sapiens or Mus musculus reference genome, version refdata-gex-GRCh38-2020-A and GRCm38-mm10 10X Genomics, respectively.

#### Quality Control and normalization

We filtered cells with less than 100 detected genes and with more than 10% of mitochondrial genes to eliminate low quality cells. Then, we estimated the number of doublet sample by sample based on the number of cells and deleted the percentage of estimated cells with higher number of genes. After this, we followed the standard workflow proposed by Seurat package^39^, normalized counts with the default “LogNormalize” method, selected the highly variable features and scaled them sample by sample.

#### Dimensionality reduction and clustering

We analyzed multiple PDTTCs in an integrated approach and paired control and treated cultured samples. Then, we performed a linear dimensionality reduction step, Principal Component Analysis (PCA), followed by an integration step using Harmony method (Harmony v1.2.0)^40^ and a clustering step using 20 dimensions of reductions to build a K-nearest neighbours graph and Louvain algorithm. Finally, we run the Uniform Manifold Approximation and Projection (UMAP) algorithm to visualize the data.

#### Cell type annotation

For identifying cell types, we first used SingleR package^41^ using the Primary Cell Atlas^42^ as reference. Then, we explored gene expression levels of markers defined in bibliography in the different clusters and checked the identified markers of each cluster with FindAllMarkers function from Seurat package. For identification of malignant cells, we also infered Copy Number Variation (CNV) with infercnv package^43^ using myeloid and lymphoid cells as reference. We used gene order file provided by TrinityCTAT corresponding to hg38 or mm10/Gencode. After cell type annotation, myeloid compartment was reclustered following the same steps described in the previous section.

#### Differential expression

Differential expression analysis was performed to explore the transcriptomic effect of anti-LIF treatment in the different cell types using Wilcoxon Rank Sum test and Bonferroni method for p-value adjustment. The anti-LIF signature was defined as the intersection of genes consistently downregulated (log2FC < -0.2 and adjusted p-value < 0.05) in the myeloid compartment in response to LIF blockade in patients 4, 6, 7 and 9. The C1q^+^ TAMs signature was derived from the integrated object comprising all nine patients across both conditions. Myeloid cells were subset from this object, and reclustering was performed within the control condition. Among the resulting clusters, clusters 0, 1 and 2 exhibited the highest expression of C1q. Marker genes distinguishing these clusters from the remaining myeloid clusters were identified using FindMarkers. The top 31 markers with an adjusted p-value of 0 were selected to define the C1q^+^ TAMs signature.

### Spatial transcriptomics (Xenium)

Spatial Transcriptomics data from PDTTCs was generated on the Xenium In Situ platform using the pre-designed Xenium Human Breast panel targeting 280 genes (chemistry version 1, cat #: 1000599). PDTTC samples were prepared according to the manufacturer’s instructions (CG000578 – Rev C) after assessing sample quality by hematoxylin and eosin staining in adjacent tissue sections from the same blocks. Two formalin-fixed paraffin-embedded sections (5 microns thick) were mounted onto a Xenium slide, incubated at 42°C for 3 hours and dried overnight at room temperature in a desiccator. The Xenium slide was processed and analyzed 4 days after sectioning. Sections were deparaffinized and de-crosslinked (CG000580 – Rev C), and then hybridized with the pre-designed probes at 50°C overnight (∼20 hours), followed by post-hybridization washes, ligation, amplification, multimodal cell segmentation staining and autofluorescence-quenching as described in user guide CG000749 – Rev A. The Xenium slide was loaded on the Xenium Analyzer instrument for imaging and analysis under software version 1.3.3.0, following the Imaging user guide CG000584 – Rev E. The downstream analysis was performed with the Seurat package. In order to remove low quality cells, cells with less than 30 detected transcripts were filtered out and the SCTransform function was used for counts normalization. The 30 principal components from PCA were used to cluster the cells with the functions FindNeighbors and FindClusters (resolution 0.3). Clusters were annotated as “Malignant” based on the expression of the gene markers *PDGFRA*, *CCND1* and *EGFR*. Endothelial, Myeloid and Lymphoid cells were annotated based mainly on the expression of the gene markers *PECAM1*, *CD163* and *CD3E*, respectively. To show the annotated segmented cells together with the location of selected molecules, the ImageDimPlot function of the Seurat package was used.

To test differences between myeloid cells of control vs anti-LIF conditions, cells annotated as Myeloid of each condition were first subset and the preprocessing steps mentioned above were repeated (normalization and PCA) to then integrate them with the Harmony R package (v.0.1.0)^40^ using the first 30 PCA dimensions.

Gene expression level of the genes “*CCR7*”, “*C1QA*” and “*MRC1*” of each condition were plotted with the ViolinPlot function. Lastly, differentially expressed genes between the conditions were tested using the FindMarkers function and the significant genes (|log_2_FC|>0.3 and adjusted p-value< 0.01) were plotted as a volcano plot.

### Generation and processing of bulk RNAseq data

Total RNA extracted from GBM tumors, PDTTCs or mouse BMDMs were quality-checked using an Agilent 2100 Bioanalyzer, and only samples with RNA Integrity Number (RIN) >7 were included. Sequencing libraries were prepared using a poly(A) enrichment protocol and sequenced on an Illumina NovaSeq 6000 platform, generating 150 bp paired-end reads with a depth of 30 million reads per sample at Centro Nacional de Análisis Genómico (GBM tumors and PDTTCs) or HaploX (mouse BMDMs). RNAseq data were pre-processed with the nf-core/rnaseq pipeline (v.3.14.0)^44^ using GRCh38 as reference genome^45^. Log_2_ transformed centered and scaled transcripts per million (TPM) normalized counts, were used to generate heatmaps with the anti-LIF signature using the ComplexHeatmap package^46^. Samples were clustered in 2 groups applying K-means clustering based on Euclidean distance. For enrichment of anti-LIF gene signature in response to anti-LIF treatment, we inferred a score through single sample gene set enrichment (ssGSEA) ^47^. For mouse BMDMs, GRCm38/mm10 was used as a Mus musculus reference genome. Differential expression analysis was conducted using DESeq2 (v1.36.0) in R. Genes with an adjusted p-value<0.05 and |log2 FC|>1 were considered significantly differentially expressed and were represented using the ComplexHeatmap package. Genes were clustered using hierarchical clustering (method: ward.D2, distance: euclidean).

### Transcriptomic analysis of clinical data sets

RNA-seq gene expression, whole-exome sequencing (WES), and clinical data from the GBM_CPTAC cohort were retrieved from the GDC Data Portal (https://portal.gdc.cancer.gov/projects/CPTAC-3)^32^. Patients harboring mutations in the IDH1 gene were excluded from all downstream analyses, including clustering and correlation analyses. To assess the association between C1q⁺ TAMs and cytokine expression, TPM-normalized RNA-seq counts were log2-transformed, centered, and scaled, and used to infer a C1q⁺ TAM signature score by single-sample gene set enrichment analysis (ssGSEA)^47^ across all patients. Partial correlations between the C1q⁺ TAM ssGSEA score and the log2-TPM expression of each cytokine were computed using the pcor.test function from the ppcor R package^48^. CD68 log2-TPM expression was included as a covariate in the partial correlation analysis as a proxy for overall myeloid/macrophage infiltration. Results were visualized as a volcano plot. Log2-TPM-normalized counts were used to generate heatmaps of the anti-LIF gene signature using the ComplexHeatmap package^46^. Patients were clustered into two groups using k-means clustering based on Euclidean distance. GBM molecular subtypes were assigned according to^32^. Gene set enrichment was performed using GSEA^49^ against HALLMARK gene sets from MSigDB, and normalized enrichment scores (NES) and p values were obtained for each patient group. Finally, the association between LIF expression and the C1q⁺ TAM signature score was evaluated by correlation analysis.

## Data availability

The analyses were performed using R4.1.2 and Seurat package v5.1. The raw transcriptomic data generated in this study have been deposited in the GEO database repository under accession code GSE304807 and GSE330393, or at the European Genome-phenome Archive (EGA), under accession number EGAS50000001810.

## Author contributions

J.S. designed this study. M.L.V., S.E.G., R.I. and L.C.B. contributed to the design and development of the experiment. M.L.V., S.E.G., R.I., L.C.B., L.C.C, E.B.T. and A.A. performed or supported *in vitro* experiments. M.L.V., S.E.G., R.I., L.C.B., A.G.E, D.S.S., L.C.C., C.G.M., I.C. and M.D. performed or supported *in vivo* experiments. A.N.A, C.M.A. processed and analysed RNA-seq and scRNA-seq data. I.S. and H.H. processed and analysed the spatial transcriptomic data. P.C., M.G.A. and S.G. provided technical support on the development of CAR T-cells. J.G., E.P. and F.M.R. participated in patient sample and clinical data collection.

## Acknowledgements

This study was undertaken with the support of ISCIII, Immune4all (PMP22/00054, R.I.); Fundación Asociación Española contra el Cáncer (AECC); ISCIII, FIS (PI22/00130); PRE2021-096899 (M.L.V.); PRE2022-102567 (L.C.B.); LCF/BQ/DR22/11950016 (C.G.M.); JDC2023-052465-I (S.E.G.) and PRDBA2610350SOLI (D.S.S.); LCF/BQ/DFR25/12000071 (C.M.A.).

## Supplementary

**Supplementary Fig. 1.**
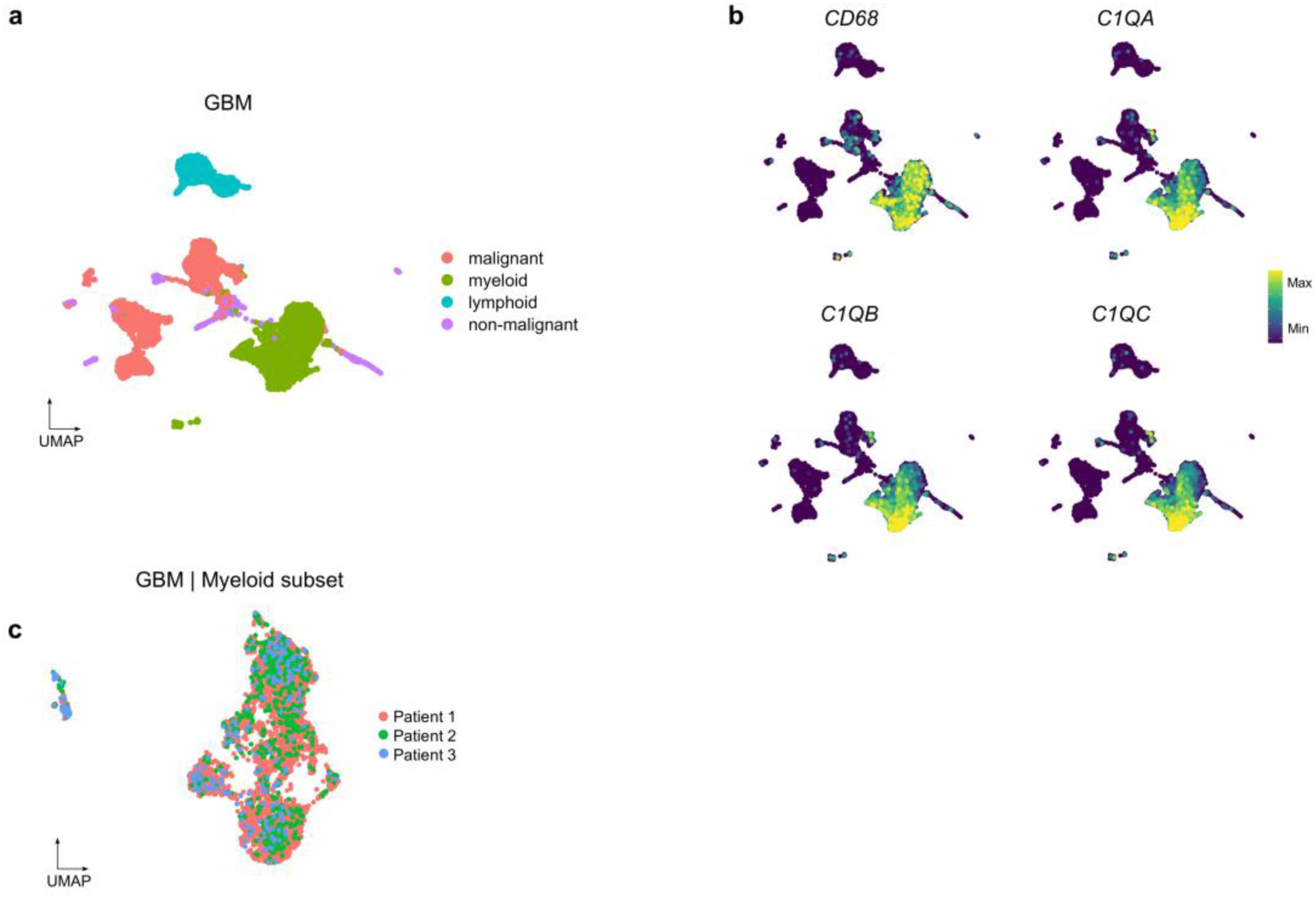
Myeloid cells expressing C1q in GBM, related to Fig. 1. **a,** UMAP of scRNAseq data from 3 GBM patients from (Iurlaro et al. co-submitted). **b**, UMAP coloured according to gene expression levels of *CD68*, *C1QA*, *C1QB* and *C1QC*. **c**, UMAP of the myeloid cell subset clustered by patients.

**Supplementary Fig. 2.**
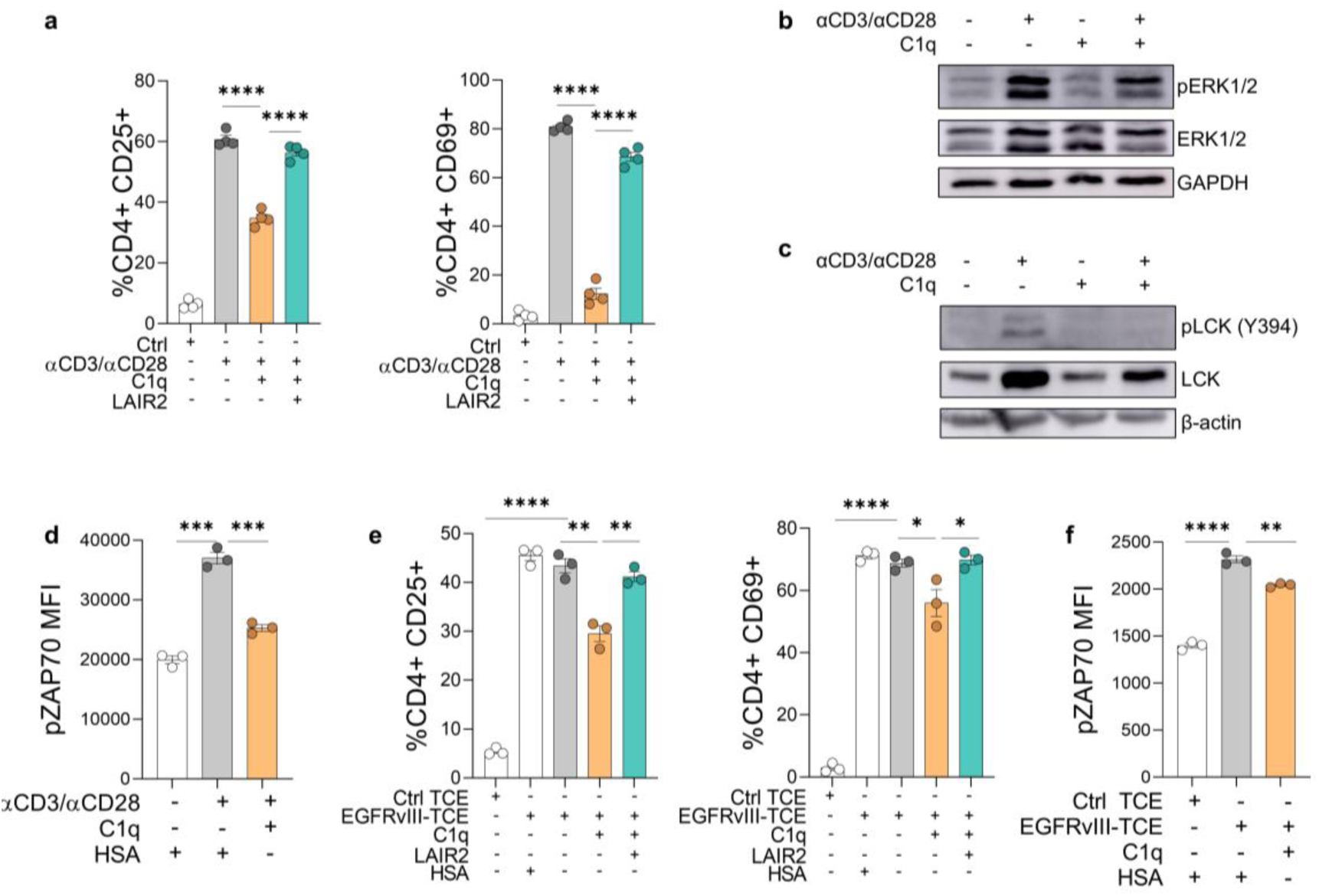
C1q impairs T cell receptor associated signaling, related to Fig. 2. **a,** T-cell activation in PBMCs pre-incubated overnight with 100 μg/mL C1q or HSA with or without LAIR2 and activated for 24h with soluble αCD3/αCD28. T-cell activation was measured as percentages of CD25^+^ and CD69^+^ cells in CD4^+^ T-cells (n=4). **b**, Immunoblot of pERK1/2 and ERK1/2 in human PBMCs. When indicated, cells were pre-incubated with 100 μg/mL C1q overnight and stimulated for 15min with soluble αCD3/αCD28. GAPDH was used as a loading control. **c**, Immunoblot of pLCK (Y394) and LCK in human CD3^+^ T-cells. When indicated, cells were pre-incubated with 100 μg/mL C1q overnight and stimulated for 15min with soluble αCD3/αCD28. β-actin was used as a loading control. **d**, TCR activation in Jurkat cells pre-incubated overnight with 100 μg/mL C1q or HSA and then activated with soluble αCD3/αCD28 for 3min at 37°C. TCR activation was measured as mean fluorescence intensity of pZAP70 (n=3). **e**, T-cell activation in PBMCs pre-incubated for 2h with 100 μg/mL C1q or HSA and then co-cultured with U87-EGFRvIII cells. When indicated, C1q and LAIR2 were added to the co-culture prior to the addition of EGFRvIII-TCE or control TCE. T-cell cytotoxicity was measured as tumor cell death by flow cytometry. T-cell activation was measured as percentages of CD25^+^ and CD69^+^ cells in CD4^+^ T-cells (n=3). **f,** TCR activation in CD3^+^ T-cells pre-incubated overnight with 100 μg/mL C1q or HSA and then co-cultured with U87-EGFRvIII cells for 1h in the presence of EGFRvIII-TCE or control TCE. TCR activation was measured as mean fluorescence intensity of pZAP70 (n=3). Data are mean ± SEM. Statistical analyses by Student’s *t*-test (**a**), (**d**), (**e**), (**f**). \**p*<0.05; \*\**p*<0.01; \*\*\**p*<0.001; \*\*\*\**p*<0.0001.

**Supplementary Fig. 3.**
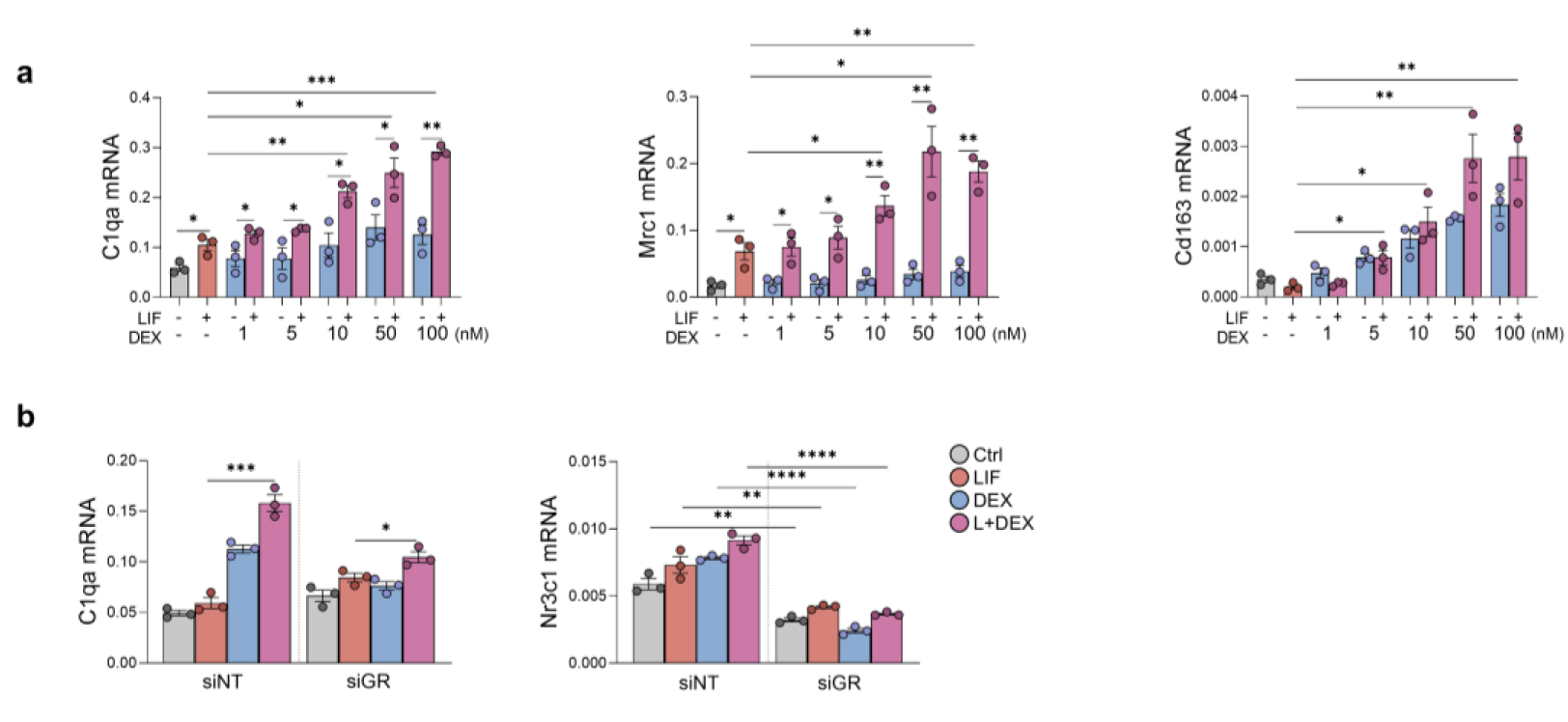
C1q regulation by LIF and dexamethasone, related to Fig. 3. **a,** mRNA expression of the indicated genes in BMDMs treated with 20 ng/mL LIF for 72h and different concentrations of dexamethasone for the last 24h (n=3). **b**, *C1qa* mRNA expression in BMDMs transfected with non-targeting siRNA or siRNA against the glucocorticoid receptor and treated with 20 ng/mL LIF for 24h and 100 nM dexamethasone for the last 24h. Comparison between LIF and LIF+DEX groups. *Nr3c1* mRNA expression used to assess the efficacy of silencing. Data are mean ± SEM. Statistical analyses by Student’s *t*-test (**a**), (**b**). \**p*<0.05; \*\**p*<0.01; \*\*\**p*<0.001; \*\*\*\**p*<0.0001.

**Supplementary Fig. 4.**
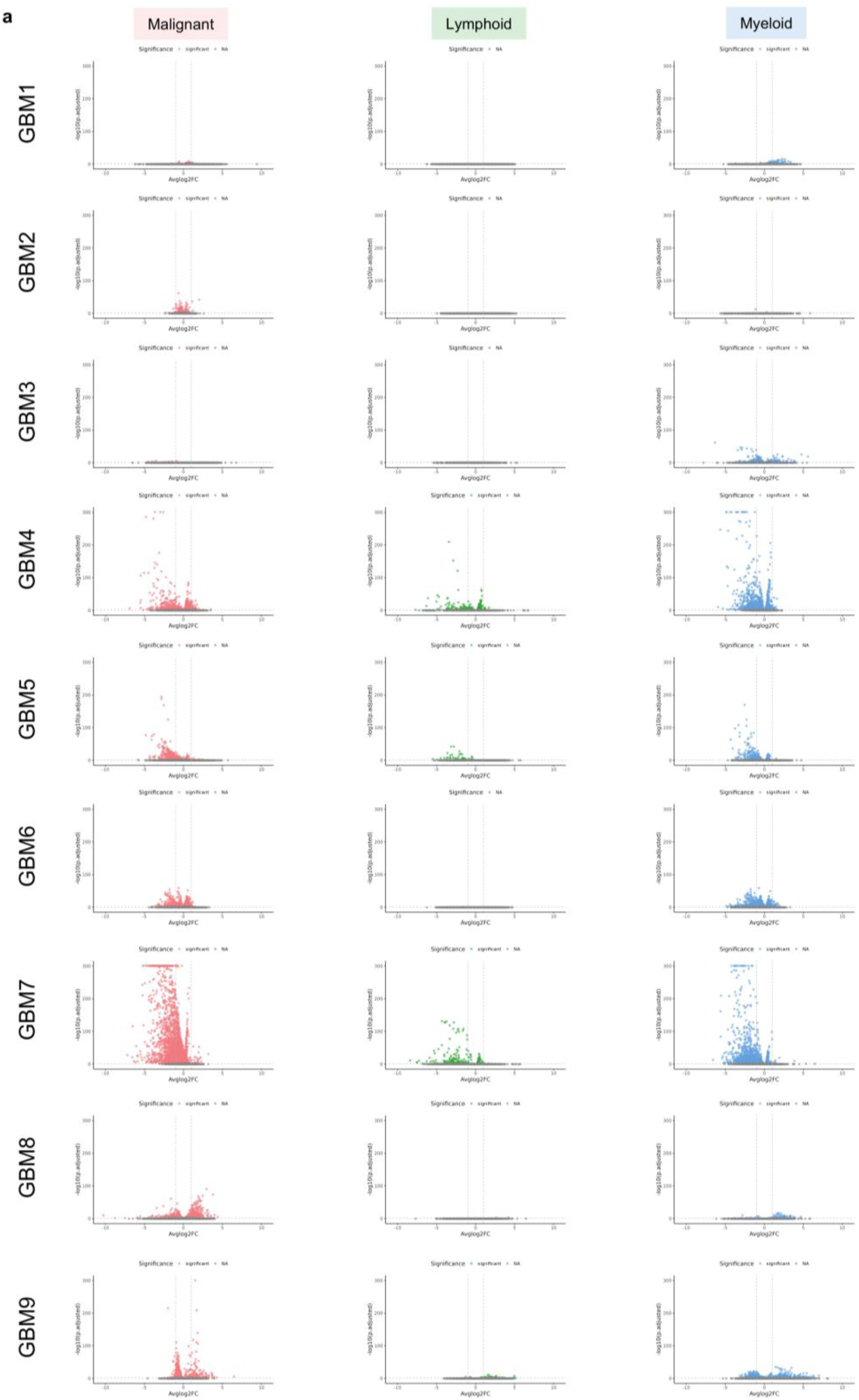
Response to LIF blockade is heterogeneous among GBM samples, related to Fig. 4. **a,** Volcano plots representing the differentially expressed genes (adjusted p-value<0.05) in the malignant (left panel), lymphoid (center panel) and myeloid populations (right panel) in response to LIF blockade for all 9 PDTTCs.

**Supplementary Fig. 5.**
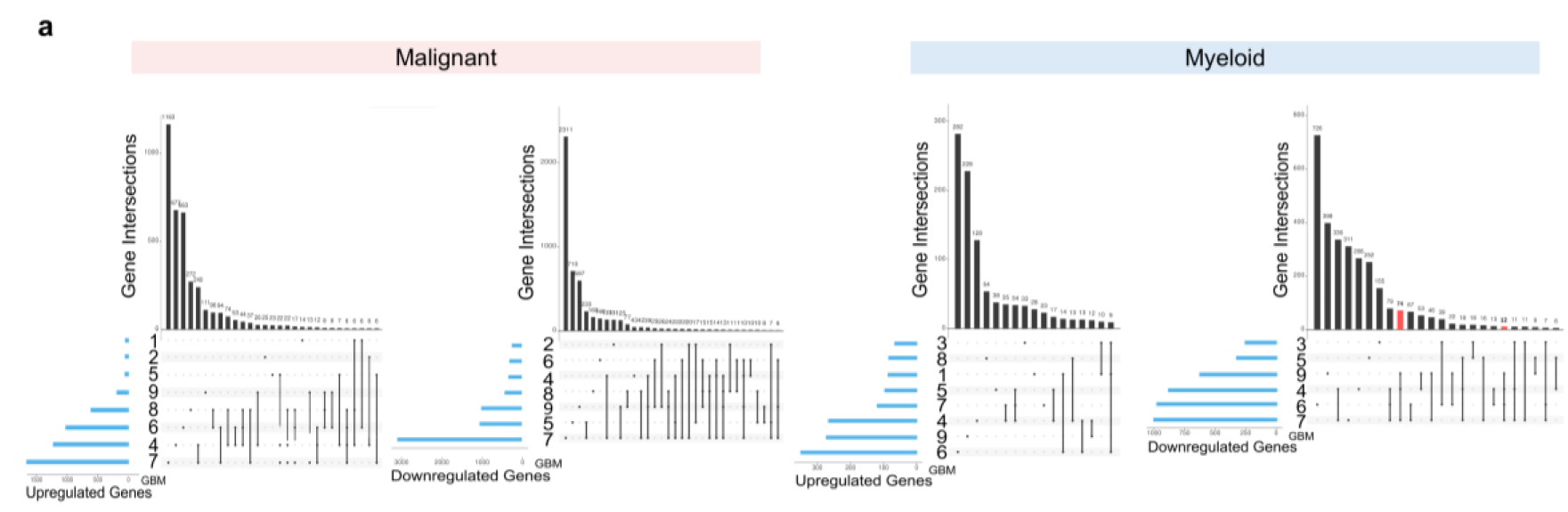
Common downregulated genes in the myeloid population define an anti-LIF signature, related to Fig. 4. **a,** UpSet plots showing the intersection of the differentially expressed genes (DEGs) in malignant (left panels) and myeloid (right panels) populations across all samples upon anti-LIF treatment. Marked in red, the 74 and 12 genes intersections that define the anti-LIF signature. Samples with no DEGs or intersections of less than 6 genes were excluded.

**Supplementary Fig. 6.**
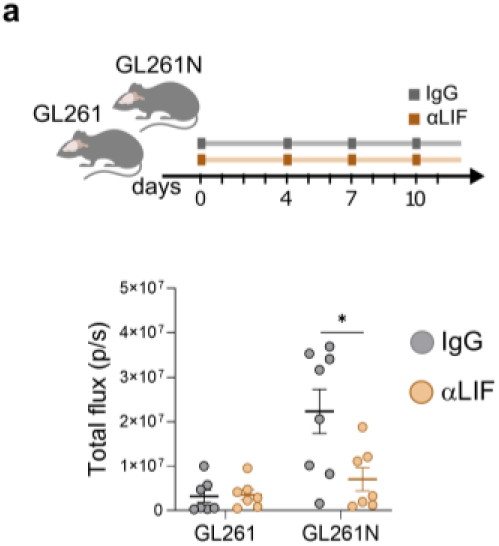
LIF blockade reduces tumor growth in GL261N orthotopic mouse model, related to Fig. 5. **a**, Top panel, schematic representation of the experiment. Mice were inoculated with either GL261 or GL261N cell lines and treated from the day of surgery with IgG or anti-LIF. Tumor growth upon anti-LIF treatment assessed by bioluminescence imaging (n=7 GL261 IgG, n=7 GL261 anti-LIF, n=8 GL261N IgG, n=7 GL261N anti-LIF). Data are represented as mean ± SEM. Statistical analyses by Mann–Whitney t-test (a). \**p*<0.05.

**Supplementary Fig. 7.**
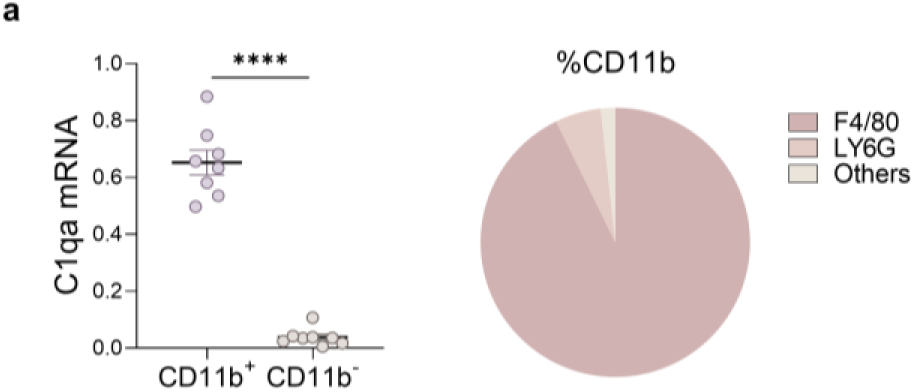
C1q expression is restricted to myeloid cells, related to Fig. 5. **a**, Left panel, *C1qa* expression in the positive fraction (CD11b^+^) and negative fraction (CD11b^-^) obtained after CD11b^+^ cells isolation from GL261N tumors (n=8). Right panel, myeloid populations in isolated CD11b^+^ cells from GL261N tumors (n=3). Data are mean ± SEM. Statistical analyses by Student’s *t*-test.**** *p*<0.0001.

